# A heptavalent O-antigen bioconjugate vaccine exhibits differential functional antibody responses against diverse *Klebsiella pneumoniae* isolates

**DOI:** 10.1101/2023.12.12.571344

**Authors:** Paeton L. Wantuch, Cory J. Knoot, Lloyd S. Robinson, Evgeny Vinogradov, Nichollas E. Scott, Christian M. Harding, David A. Rosen

## Abstract

*Klebsiella pneumoniae* is a concerning pathogen that is now the leading cause of neonatal sepsis and is increasingly difficult to treat due to heightened antibiotic resistance. Thus, there is an urgent need for preventive and effective immunotherapies targeting *K. pneumoniae*. Vaccination represents a tractable approach to combat this resistant bacterium in some settings; however, there is currently not a licensed *K. pneumoniae* vaccine available. *K. pneumoniae* surface polysaccharides, including the terminal O-antigen polysaccharides of lipopolysaccharide, have long been attractive candidates for vaccine inclusion. Herein we describe the generation of a bioconjugate vaccine targeting seven of the predominant O-antigen subtypes in *K. pneumoniae*. Each of the seven bioconjugates were immunogenic in isolation, with limited cross-reactivity among subtypes. Vaccine-induced antibodies demonstrated varying degrees of binding to a wide variety of *K. pneumoniae* strains, including suspected hypervirulent strains, all expressing different O-antigen and capsular polysaccharide combinations. Further, sera from vaccinated mice induced complement-mediated killing of many of these *K. pneumoniae* strains. Finally, we found that increased quantity of capsule interferes with O-antigen antibodies’ ability to bind and mediate killing of some *K. pneumoniae* strains, including those carrying hypervirulence-associated genes. Taken together, these data indicate that this novel heptavalent O-antigen bioconjugate vaccine formulation exhibits promising efficacy against some, but not all, *K. pneumoniae* isolates.

## Introduction

*Klebsiella pneumoniae* is a Gram-negative bacterium that frequently causes pneumonia, urinary tract infection, sepsis, and other hospital-acquired infections (1). It most often infects the immunocompromised, elderly, and very young (2, 3). Recently, *K. pneumoniae* was determined to be the most common causative agent of infectious deaths in children under the age of five in low and middle-income countries, often associated with neonatal sepsis (4, 5). Two circulating pathotypes of *K. pneumoniae* are currently recognized: classical (c*Kp*) and hypervirulent (hv*Kp*). c*Kp* isolates are typically associated with hospital-acquired infections, while hv*Kp* often causes community-acquired infections in otherwise healthy hosts (6). The recent emergence of antibiotic resistance among *K. pneumoniae* strains has caused great concern, as isolates acquire such resistance mechanisms as extended-spectrum β-lactamases (ESBLs) and *K. pneumoniae* carbapenemases (KPCs) and threaten a post-antibiotic era (7, 8). ESBL and KPC isolates have been assigned the highest levels of concern by the US Centers for Disease Control and Prevention (CDC) and World Health Organization (WHO) (9). Immunization may prove a valuable strategy to alleviate the burden of such isolates; however, no licensed vaccine currently exists for *K. pneumoniae*.

Surface polysaccharides have drawn great interest as potential targets for a *K. pneumoniae* conjugate vaccine. *K. pneumoniae* expresses two major surface polysaccharides, capsule (CPS) and the O-antigen of lipopolysaccharide (LPS) (10). While capsule may represent an efficacious vaccine target, *K. pneumoniae* as a species expresses over 100 different capsular types (11), and it has been estimated that an effective capsule vaccine for *K. pneumoniae* would require at least 24 different capsule antigens (12). Alternatively, there are only 11 known serogroups of O-antigens expressed by *K. pneumoniae,* and four of these (O1, O2, O3, and O5) account for greater than 80% of clinical isolates worldwide (12–14). Recently, structural subtypes have been discovered within the O1, O2, and O3 serogroups that are often associated with clinically important isolates (15–18). The nomenclature for these subtypes continues to evolve, both in name of the loci required for O-antigen production (e.g. O1/O2v1) and in name of the functional serotype produced (e.g. O2a) (19). For simplicity herein, we refer to the O1 and O2 O-antigen subtypes by their original Kaptive locus names for both their genotype and confirmed phenotypic serotypes.

Polysaccharides are generally poor immunogens in isolation, often failing to induce an adaptive immune response and durable immunological memory (20). This can be overcome by attachment of the polysaccharide to a carrier protein, termed conjugation. Conjugate vaccines have demonstrated great success in targeting other bacterial pathogens including *Streptococcus pneumoniae*, *Haemophilus influenzae*, and *Neisseria meningitidis*, and therefore have been an area of focus for *K. pneumoniae* vaccine development. A unique method of polysaccharide protein conjugation, termed bioconjugation, has previously been applied to produce *K. pneumoniae* polysaccharide-protein conjugates that feature an unaltered polysaccharide antigen (21, 22). Our bioconjugation technology synthesizes the polysaccharide-protein conjugate entirely within a glycoengineered bacterial expression system, allowing for rapid production of diverse polysaccharide protein conjugates targeting either capsule or O-antigen (23).

Herein we describe the development and preclinical evaluation of a broad-spectrum, heptavalent bioconjugate vaccine targeting the O1v1, O1v2, O2v1, O2v2, O3, O3b, and O5 O-antigen subtypes of *K. pneumoniae*. We synthesized these bioconjugate vaccines using our streamlined glycoengineered bacterial expression system. Purified bioconjugate vaccines were then tested for immunogenicity in CD-1 outbred male and female mice. Vaccine-induced anti-O-antigen antisera were tested for binding and complement-mediated bacterial killing against a wide variety of clinical *K. pneumoniae* isolates. Capsule quantification indicated that the amount of capsular polysaccharide produced by various strains affects antibody binding and function. These heptavalent bioconjugate O-antigen vaccine data provide critical functional vaccine information that must be considered in targeting this increasingly resistant pathogen.

## Results

### Heterologous expression of O-antigen polysaccharides for bioconjugate vaccine production

In order to create a broad vaccine targeting the majority of clinically relevant *K. pneumoniae* isolates, we sought to produce bioconjugates incorporating the seven most common O-antigen subtypes: O1v1 (O1), O1v2 (O1afg), O2v1 (O2a), O2v2 (O2afg), O3, O3b, and O5 (Figure 1). We cloned the *K. pneumoniae* O-antigen gene clusters encoding for the biosynthetic machinery required for producing each polysaccharide antigen and subsequently expressed each *in trans* in *Escherichia coli*. Structurally, *K. pneumoniae* O3 and O5 are identical to *E. coli* O9 and O8 O-antigens, respectively (24) and were cloned directly from reference *E. coli* O9 and O8 strains. These cloned genes were introduced into self-replicating vector backbones and expressed in combination for heterologous expression of *K. pneumoniae* O-antigens in *E. coli*. Six of the *K. pneumoniae* O-antigen clusters were produced in *E. coli* (O1v1, O2v1, O2v2, O3, O3b, and O5); nuclear magnetic resonance spectroscopy analysis of extracted polysaccharides from *E. coli* confirmed structural matches to the native *K. pneumoniae* O-antigen polysaccharides, including terminal caps on the O3, O3b, and O5 polysaccharides (Supplemental Figures 1-5). The NMR structure of the O1v1 bioconjugate was previously reported (21). The O1v2 bioconjugate was produced directly in a genetically modified *K. pneumoniae* NTUH-K2044 strain (hereafter referred to as NTUH), which naturally produces the O1v2 O-antigen polysaccharide. To enhance O1v2 bioconjugate production in subsequent experiments, the *wcaJ* and *waaL* genes were deleted from the NTUH chromosome. The *wcaJ* deletion prevents the synthesis of K1 capsule, which might otherwise be transferred to the carrier protein and contaminate the O1v2 bioconjugate. The *waaL* deletion prevents the nascent O-antigen polysaccharide precursor from being transferred to the *Klebsiella* core oligosaccharide. The resulting strain was then transformed with a plasmid encoding the carrier protein and oligosaccharyltransferase (OTase). The carrier protein used for all *K. pneumoniae* bioconjugates was a genetically inactivated *Pseudomonas aeruginosa* exotoxin A (EPA) engineered to contain two internal 23-amino acid sequons that are glycosylated by the *Acinetobacter baylyi* ADP1 PglS OTase (21, 25).To produce the bioconjugates, EPA and PglS were co-expressed from plasmid pVNM245 (21) alongside the *K. pneumoniae* O-antigen expression plasmids in *E. coli* strain CLM24 (26) or *K. pneumoniae* NTUH*ΔwcaJΔwaaL*. Bacteria were cultured in Terrific Broth and induced with 0.1 mM IPTG at mid-log followed by overnight growth. The seven bioconjugates and unglycosylated EPA carrier were purified from frozen cell pellets using a combination of Ni-affinity chromatography, anionic exchange chromatography, and size exclusion chromatography.

**Figure 1.**
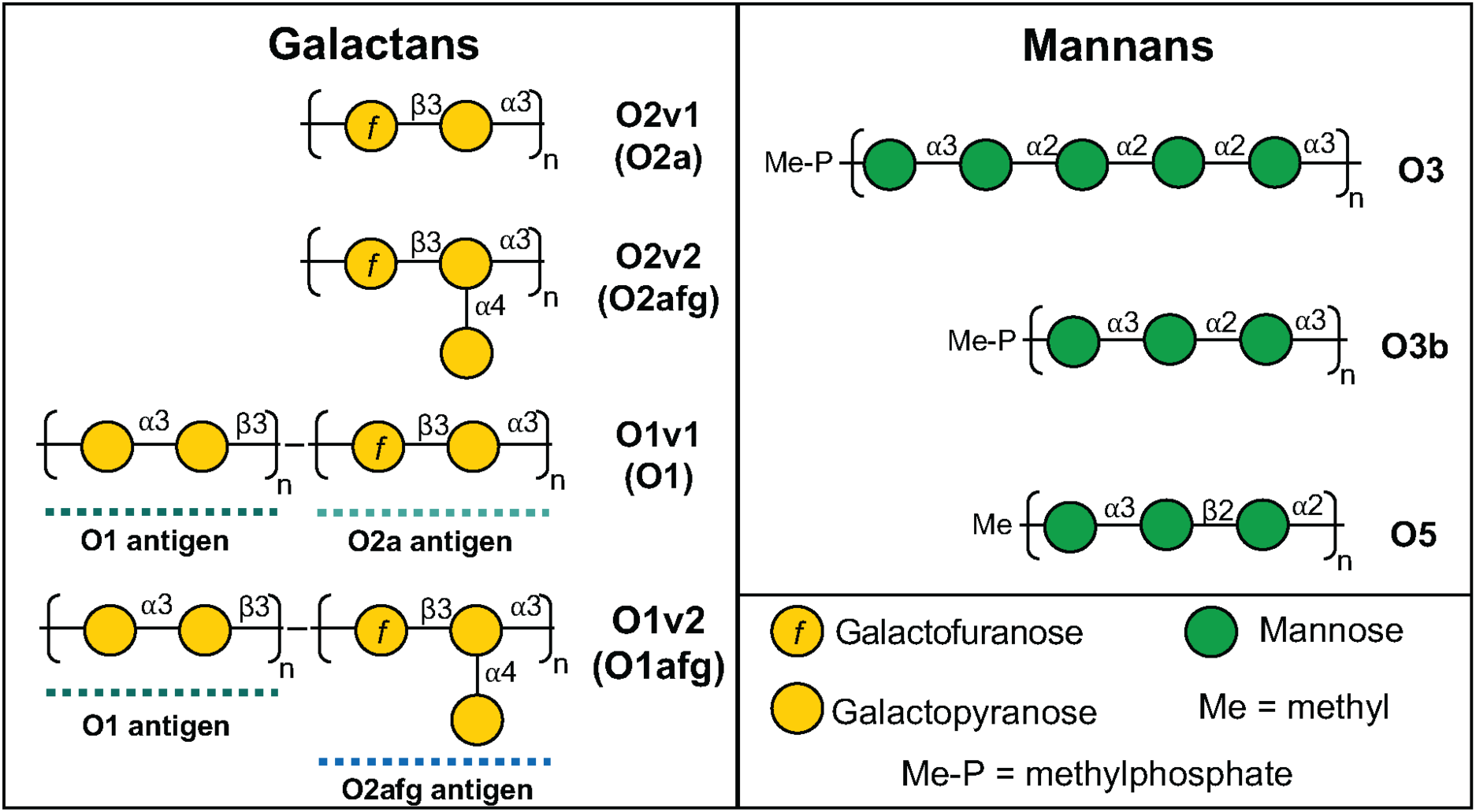
Structures of the seven most common *K. pneumoniae* O-antigens subtypes.

### Characterization of O-antigen bioconjugate vaccines

Purified O-antigen-EPA bioconjugates were first characterized via Coomassie staining and western blotting followed by intact mass spectrometry prior to murine studies. Each biconjugate was probed with anti-6x-His protein antibody (to recognize tagged EPA) and separately with primary antibodies derived from mouse or rabbit *K. pneumoniae* O-antigen antisera (Figure 2). O2v1 mouse antisera showed cross-reactivity with O1v1 due to the presence of the conserved O2a antigenic structure in the O1v1 antigen (Figure 2A – C). Interestingly, the O2v1 antisera also co-reacted with the O2v2-bioconjugate indicating that a fraction of the O2v2-bioconjugate may consist of a fractional subset of O2a antigen. As expected, O2v2 mouse antisera were reactive towards both O2v2– and O1v2-bioconjugates which both share the O2afg antigen (Figure 2D – F). O1v1 rabbit antisera recognized both O1v1– and O1v2-bioconjugates from *E. coli* and *K. pneumoniae*, respectively (Figure 2G – I), again likely due to the presence of the common O1 antigen. Some cross-reactivity was seen between the O3 mouse antisera and the O3– and O3b-bioconjugates (Figure 2J – L), while the O5 mouse antisera was only specific for the corresponding bioconjugate (Figure 2M – O). Importantly, intact mass spectrometry of the O1v1, O2v1, O2v2, O3, O3b, and O5 bioconjugates showed the EPA carrier protein heterologously glycosylated with multiple different glycoforms, consistent with the canonical modal expression of the known O-antigen polysaccharide structures (Supplemental Figures 6-11).

**Figure 2.**
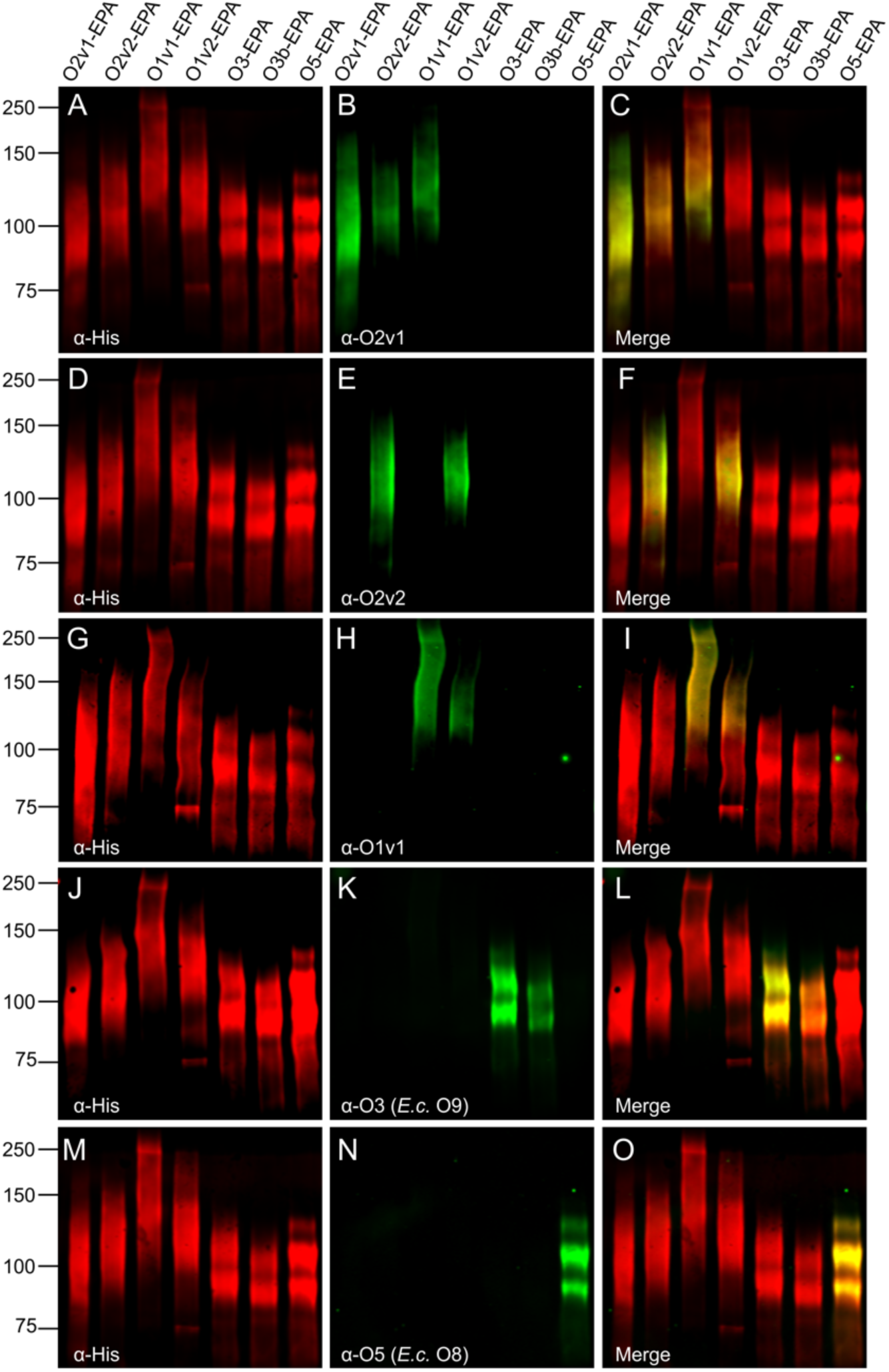
Individual western blot characterization of seven purified *K. pneumoniae* O-antigen bioconjugates probed with anti-EPA carrier protein antibody and rabbit or mouse glycan antisera. Purified bioconjugates were quantified and equivalent amounts (0.6 μg protein) loaded per lane. Protein mass markers in kDa are indicated next to each row. The purified bioconjugate is labeled at the top of each lane. A) Anti-His channel. B) O2v1 antisera channel. C) Merge of panels A and B. D) Anti-His channel. E) O2v2 antisera channel. F) Merge of panels D and E. G) Anti-His channel. H) O1v1 antisera channel. I) Merge of panels G and H. J) Anti-His channel. K) O3 (*E. coli* O9) antisera channel. L) Merge of panels J and K. M) Anti-His channel. N) O5 (*E. coli* O8) antisera channel. O) Merge of panels M and N.

### Monovalent and heptavalent O-antigen bioconjugates are immunogenic in mice

To test the immunogencity of each of the seven O-antigen bioconjugate vaccines, we immunized female CD-1 mice in seven monovalent groups: O1v1-EPA, O1v2-EPA, O2v1-EPA, O2v2-EPA, O3-EPA, O3b-EPA, or O5-EPA. Each vaccine dose was formulated with 1 µg of polysaccharide and adjuvanted with Alhydrogel (2% aluminum hydroxide gel). Mice were immunized on days 0, 14, and 28, and sera were collected on day 42. We measured polysaccharide-specific IgG in immune sera via enzyme-linked immunosorbent assay (ELISA) on 96-well plates individually coated with bacteria engineered to express one of the seven O-antigen types on their surfaces. We observed significant levels of IgG induced by each O-antigen bioconjugate against its matching O-antigen polysaccharide (Figure 3 A-G). We were further interested in the levels of cross-reaction, if any, among the different O-antigen sera and subtypes. Thus, we tested all monovalent immunization sera against each of the seven O-antigen-expressing engineered bacteria strains. We observed slight cross-reactivity of O1v1-bioconjugate immune sera with O1v2– and O2v1-expressing bacterial strains, albeit at levels significantly less than its matching O1v1-expressing strain (Figure 3A). On the other hand, O1v2-bioconjugate immune serum reacted highly against its matched O1v2-expressing strain, but also exhibited similar binding to O2v1-expressing bacteria and only slightly lower binding to O1v1-expressing bacteria (Figure 3B). O2v1-bioconjugate immune serum reacted only with its matched O2v1-expressin bacteria, while O2v2-bioconjugate immune serum reacted with both its cognate O 2v2-expressing bacteria O2v1-expressing bacteria (Figure 3C,D). Similarly, O3-bioconjugate immune serum reacted only with its target antigen, but O3b-bioconjugate immune serum demonstrated slight cross-reactivity with O3-expressing bacteria, albeit significantly lower than O3b antigen binding (Figure 3E,F). O5-bioconjugate immune serum reacted only with O5-expressing *E. coli*, exhibiting no cross-reactivity against other O-antigen subtypes. Taken together, these data demonstrate that each of the seven O-antigen bioconjugate vaccines is immunogenic in mice and that some subtypes within serogroups offer partial but modest cross-reactivity. In general, a second subtype within groups cross-reacted with the first, but not (or very little) vice versa (i.e., O3 does not cross-react with O3b, but O3b offers partial cross-reaction to O3); as a result, we felt it was necessary to include all seven subtypes in a multivalent vaccine mixture.

**Figure 3:**
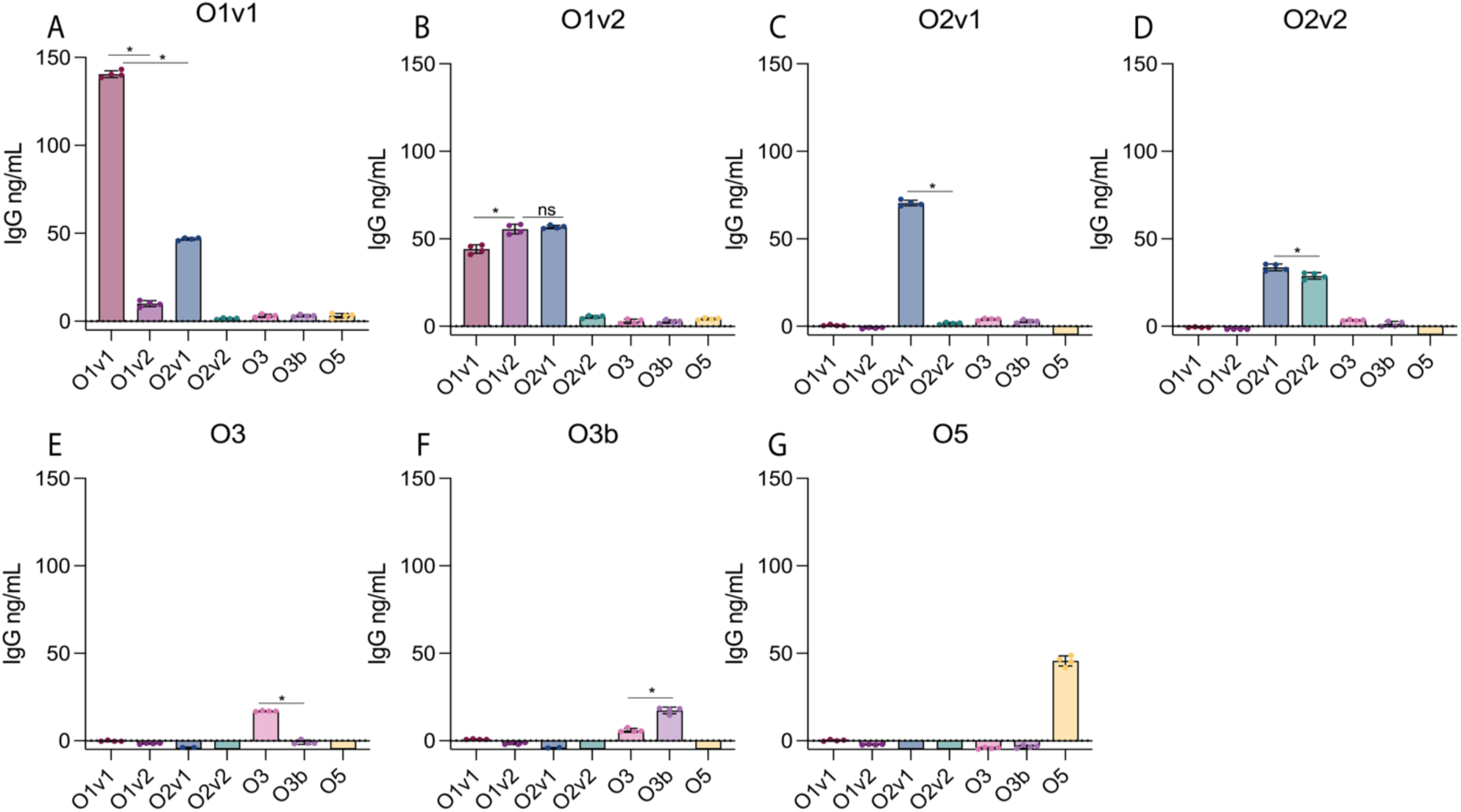
Detection of IgG generated by immunization of CD-1 mice with monovalent O-antigen bioconjugates. O-antigen-specific IgG from day 42 serum as measured in quadruplicate by ELISA using plates individually coated with engineered bacteria expressing one of the seven O-antigen subtypes. Plates were coated with bacteria expressing A) O1v1 B) O1v2 C) O2v1 D) O2v2 E) O3a F) O3b G) O5. Probing mouse sera is labeled across the x axes. Statistical analyses were performed via Mann-Whitney U test. * p<0.01; ns indicates not significant. Error bars represent means with standard deviations.

We included each of these bioconjugates (1 µg each polysaccharide per dose) to formulate a heptavalent vaccine termed O7V-EPA. To confirm that this heptavalent vaccine demonstrates similar efficacy across sexes. As such, male and female CD-1 mice were immunized as described above in two groups: control immunization (unglycosylated EPA protein) or O7V-EPA. Sera were collected on day 42 after the second booster dose. As before, polysaccharide-specific IgG was measured using ELISA on plates coated with engineered bacteria expressing individual *K. pneumoniae* O-antigens (i.e., no capsule polysaccharide expression). Compared to day 0 (pre-immune) sera, O7V-EPA-vaccinated mice exhibited significant levels of IgG against each of the seven O-antigen types (Figure 4). No significant 42-day antibody differences were observed between male and female mice (p values: 0.65 [O1v1]; 0.08 [O1v2]; 0.46 [O2v1]; 0.38 [O2v2]; 0.46 [O3]; 0.74 [O3b]; 0.86 [O5]); therefore, pooled O7V-EPA sera from both male and female mice were used for the remaining experiments.

**Figure 4:**
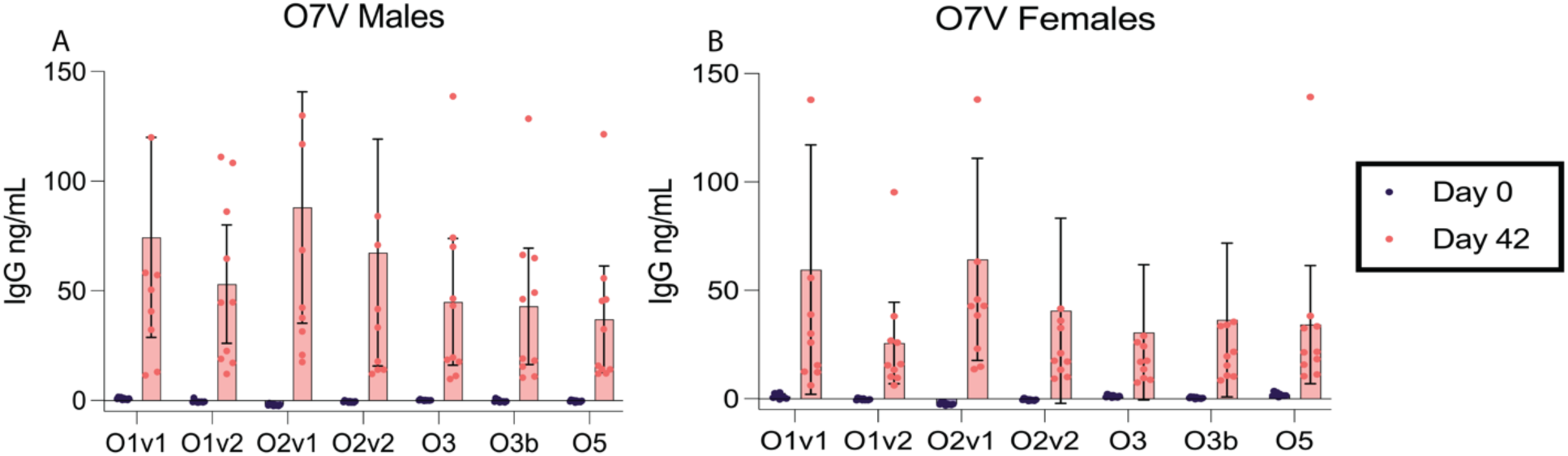
Detection of IgG generated by immunization of CD-1 male and female mice with the heptavalent O7V-EPA vaccine. O-antigen specific IgG levels were quantified at day 0 (pre-immunization) and day 42 (following full immunization series) for each mouse via ELISA with engineered bacteria expressing one of the seven O-antigen subtypes in A) male and B) female CD-1 mice. Individual CD-1 mouse IgG titers are depicted. Statistical analyses via Mann-Whitney U tests demonstrate that all day 42 sera demonstrate significant IgG titers relative to day 0 pre-immune sera. No statistical differences were detected between male and female day 42 sera. Error bars represent means with standard deviations.

### O-antigen immune sera reacts with O-matched strains of *Klebsiella pneumoniae*

We next sought to determine whether O-antigen antibodies reacted with clinical strains of *K. pneumoniae*, which produce both O-antigen containing LPS and capsular polysaccharide on their surface. We selected two strains of each O-antigen subtype from the BEI Resources Repository (27) (*K. pneumoniae* Multidrug-Resistant Organism Repository and Surveillance Network Diversity Panel) and included two known heavily-encapsulated strains 43816 and NTUH and their respective capsule knockouts (Table 1). The strains chosen encoded a range of capsule types. O-antigen-specific IgG was measured via ELISA on plates coated with each of the 18 different strains of *K. pneumoniae* using pooled sera from O7V-EPA immunized mice. O7V-bioconjugate immune serum reacted with 16 of the 18 strains to varying degrees (Figure 5). O7V-bioconjugate immune serum did not react with strains 43816 and NTUH; however, it reacted highly with 438161*cps* and NTUH1*cps* mutants, indicating that the heavy capsule production by wild-type 43816 and NTUH masked the O-antigen. Interestingly, O7V-bioconjugate immune serum reacted to differing degrees with the 14 BEI isolates, with the lowest reactivity toward strains 28183, 669448, 73059, and 1912. Generally, strains noted to encode hypervirulence-associated genes exhibited lower levels of antibody binding, except for strain 20522.

**Table 1:** Strains of *K. pneumoniae* used in this study. Multi drug-resistant and extensively drug-resistant were defined as previously described (27).

| *K. pneumoniae*<br>Strain | O-Antigen<br>Type | Capsule<br>Type | Antibiotic<br>Resistance | Virulence<br>Genes | Site of<br>Isolation |
| --- | --- | --- | --- | --- | --- |
| BEI 740795 | O1v1 | K2 | MDR | - | Respiratory |
| BEI 702261 | O1v1 | K54 | MDR | + | Wound |
| BEI 371351 | O1v2 | K57 | MDR | - | Wound |
| BEI 19073 | O1v2 | K28 | MDR | - | Wound |
| BEI 20522 | O2v1 | K24 | MDR | + | Respiratory |
| BEI 28183 | O2v1 | K154 | XDR | - | Respiratory |
| BEI 669448 | O2v2 | K102 | XDR | + | Blood |
| BEI 761403 | O2v2 | K107 | XDR | - | Blood |
| BEI 7076 | O3 | K25 | XDR | - | Wound |
| BEI 28893 | O3 | K125 | Susceptible | - | Urine |
| BEI 4759 | O3b | K38 | MDR | - | Urine |
| BEI 730567 | O3b | K46 | Susceptible | - | Blood |
| BEI 1912 | O5 | K25 | Susceptible | + | Perianal |
| BEI 591344 | O5 | K61 | MDR | - | Wound |
| 43816 | O1v1 | K2 | Susceptible | + | Unknown |
| NTUH-K2044 | O1v2 | K1 | Susceptible | + | Liver Abscess |
| 43816Δ*cps* | O1v1 | N/A | Susceptible | + | Unknown |
| NTUHΔ*cps* | O1v2 | N/A | Susceptible | + | Liver Abscess |
| **MDR = multi drug-resistant ; XDR = extensively drug-resistant** |  |  |  |  |  |

Importantly, we also tested the monovalent sera against each of the 18 strains to test for cross-reactivity (Supplemental Figure 12). We noted reactivity of each monovalent immune serum to its matched O-antigen strain but saw cross-reactivity only between O1v1 and O1v2 strains. With some strains we observed lower reactivity from monovalent sera but increased reactivity from the O7V-EPA pooled serum. This might reflect variable and lower antibody concentrations among the monovalent-immunized mice, or could indicate increased immunogenicity when all O-antigens are delivered synchronously in the heptavalent formulation. Finally, as with the O7V-EPA pooled serum, we observed no recognition of 43816 or NTUH by the relevant monovalent sera (O1v1 and O1v2 respectively); however, 438161*cps* and NTUH1*cps* were readily bound by their matched O1v1– and O1v2-bioconjugate immune sera. Together these data demonstrate that the O7V-EPA immune sera are reactive to varying degrees against 18 different strains of *K. pneumoniae* that express different O-antigen and capsule types.

### O-antigen immune sera mediate killing of many *K. pneumoniae* isolates

Having quantified the presence of O-antigen-specific antibodies, we next utilized a serum bactericidal assay to assess the capacity of O7V-EPA immune sera to evoke complement-mediated bacterial killing (Figure 6). All 18 strains of *K. pneumoniae* were individually incubated with heat-inactivated O7V-EPA pooled serum and baby rabbit complement to assess bacterial killing. We observed little to no killing (0-5%) in six of the 18 strains (702261, 28183, 6694487, 28893, 43816, and NTUH), moderate killing (6-38%) in six strains (761403, 7076, 73059, 1912, 591344, and 43816Δ*cps*), and robust killing (>50%) in the remaining six strains (Figure 6). Correlating with the ELISA studies, the loss of capsule in 43816 and NTUH enabled significantly more killing compared to wild type, from 0% killing to 38% (p=0.0008) and 50% (p=0.0143), respectively (Figure 6). Except for strain 20522, all other strains encoding hypervirulent genes exhibited low levels of bacterial killing. Additionally, we noted that strain 740795 was susceptible to complement alone, as even control wells lacking immune sera demonstrated some bacterial killing. Taken together, these data suggest that O7V-EPA immune sera is efficient at killing some, but not all, strains of *K. pneumoniae* in a complement-mediated manner.

**Figure 5:**
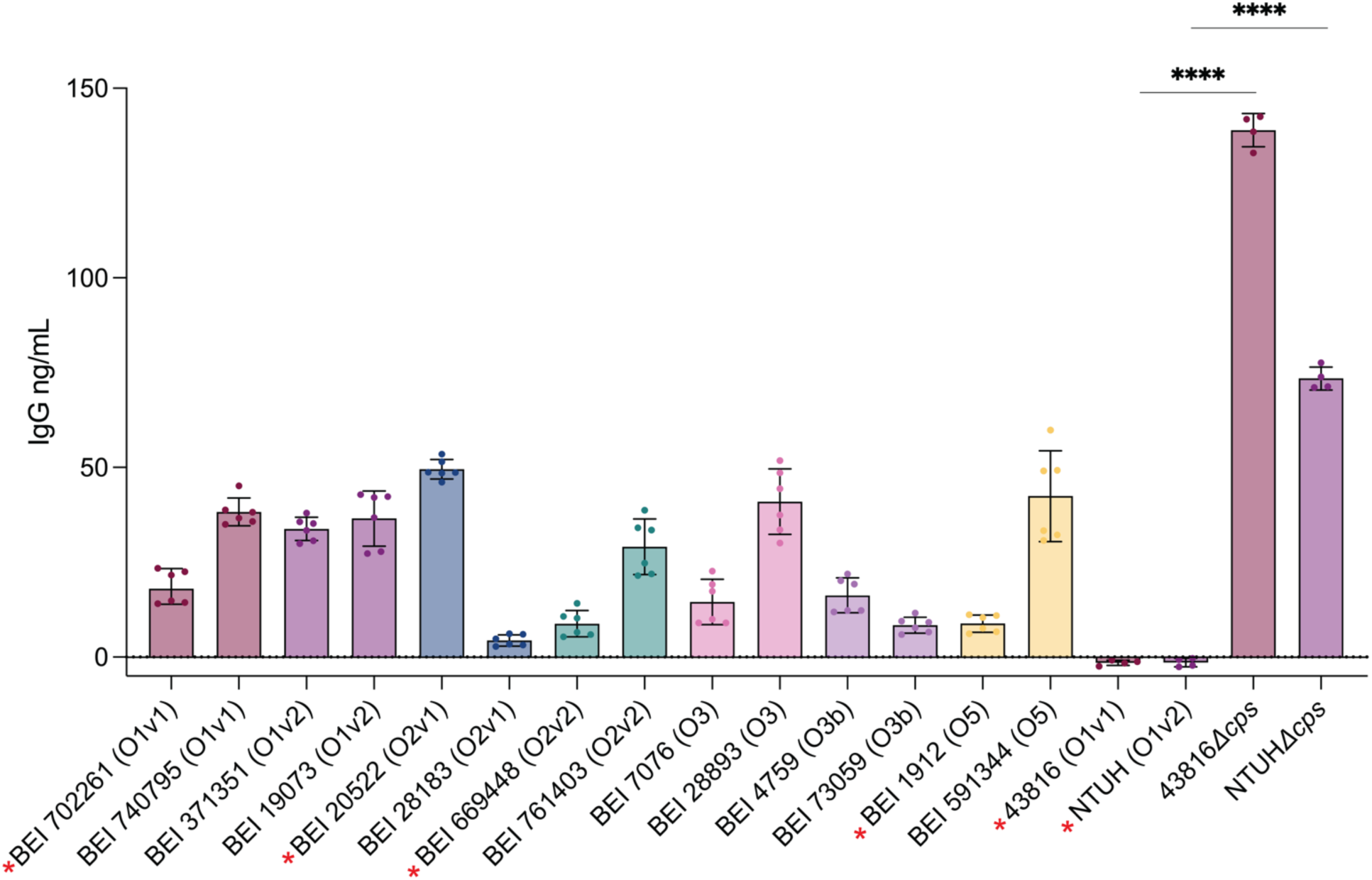
Binding of O7V-EPA bioconjugate murine vaccination-generated IgG to a panel of clinical *K. pneumoniae* isolates. O7V-EPA-generated IgGs were measured at day 42 by ELISA with plates coated with whole bacteria representing 18 diverse strains of *K. pneumoniae*. Strains with the presence of hypervirulent genes are denoted with a red asterisk. Statistical analyses were performed via Mann-Whitney U test. **** p<0.00001. Error bars represent means with standard deviations.

**Figure 6:**
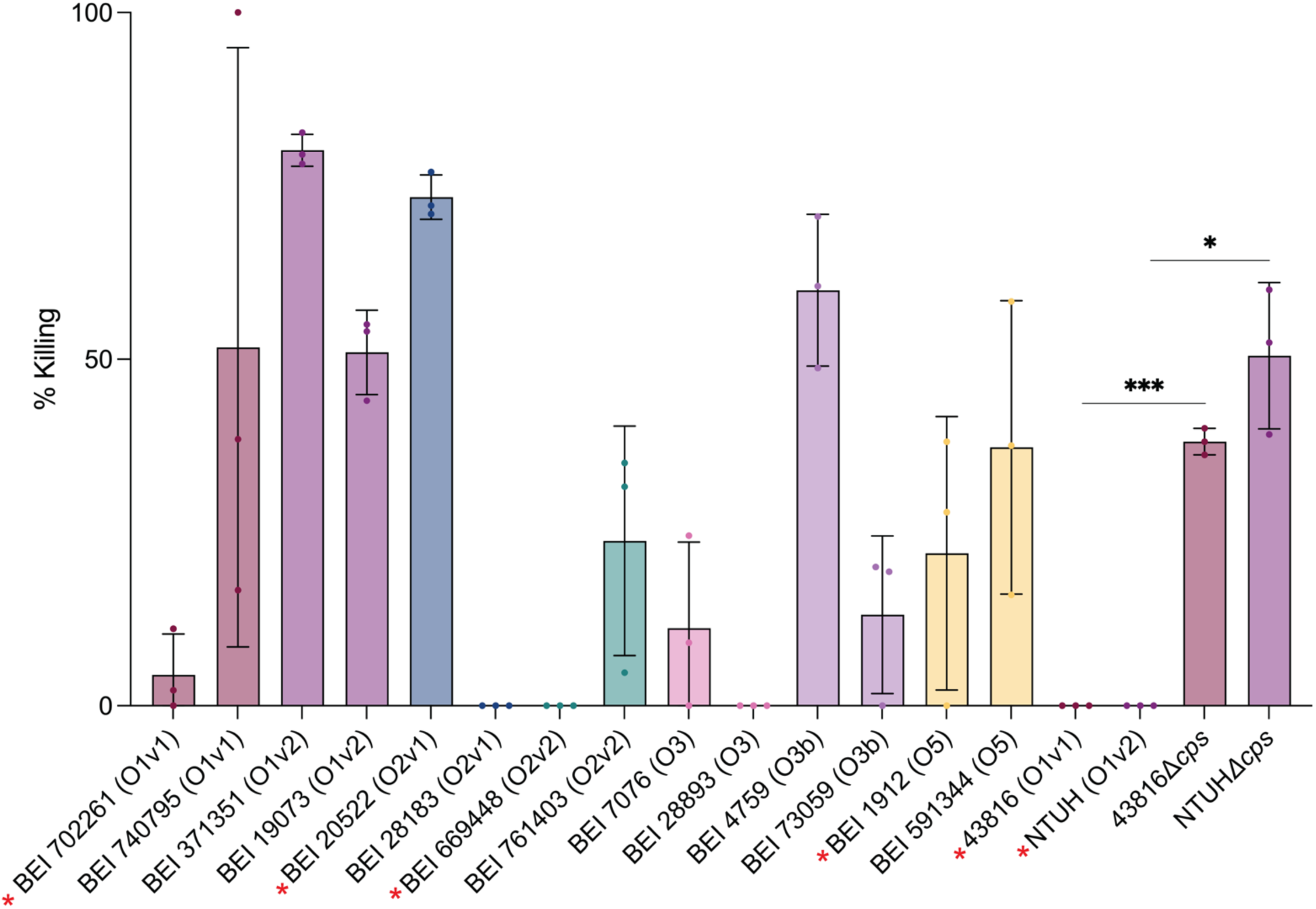
Serum bactericidal assays using heat-inactivated, pooled O7V-EPA-vaccinated mouse sera against a panel of *K. pneumoniae* clinical isolates. Serum bactericidal activities of pooled day 42 sera from mice immunized with the O7V-EPA bioconjugate were measured against 18 different strains of *K. pneumoniae*. Strains with the presence of hypervirulent genes are denoted with a red asterisk. Statistical analyses were performed via Mann-Whitney U tests. * p<0.01; *** p<0.0001; ns indicates not significant. Error bars represent means with standard deviations.

### Higher capsule levels correlate with diminished antibody binding and function

The levels of antibody binding (Figure 5) and complement-mediated killing of these strains (Figure 6) follow similar trends. Strains that demonstrated high levels of antibody reactivity also demonstrated the most bactericidal killing and vice versa; except for strain 28893, which displayed high antibody binding but no bacterial killing, and strain 4759, which displayed modest levels of antibody binding but high bacterial killing. We hypothesized that the level of capsule produced by some strains may explain the poor O-antibody binding and function against these strains, as it has been demonstrated that capsule masking of O-antigen occurs in some *K. pneumoniae* isolates (21, 28, 29). Using a glucuronic acid detection assay, we quantified the relative amount of capsule produced by each of the 18 strains. Clinical strains demonstrated vastly different amounts of capsule (Figure 7A). Negative controls (438161*cps* and NTUH1*cps*) defined the lower limit of detection. Surprisingly, three other strains (740795, 371351, and 19073) also demonstrated little to no capsule production. Interestingly, it appeared that the trend of this graph was inverse to the trend of bacterial killing and antibody response, i.e., strains making more capsule exhibited less antibody binding and functionality. There was a significant negative correlation (r^2^=0.4820; p<0.0001) between the amount of capsule present and percent killing in the serum bactericidal assay (antibody functionality) (Figure 7B). Further, with one exception (strain 20522), strains that encoded hypervirulence genes (red asterisks) exhibited the least amount of bacterial killing and had higher amounts of capsule (Figure 7B). These data indicate that strains with higher levels of capsule production exhibit decreased O-antigen antibody binding and functionality.

**Figure 7:**
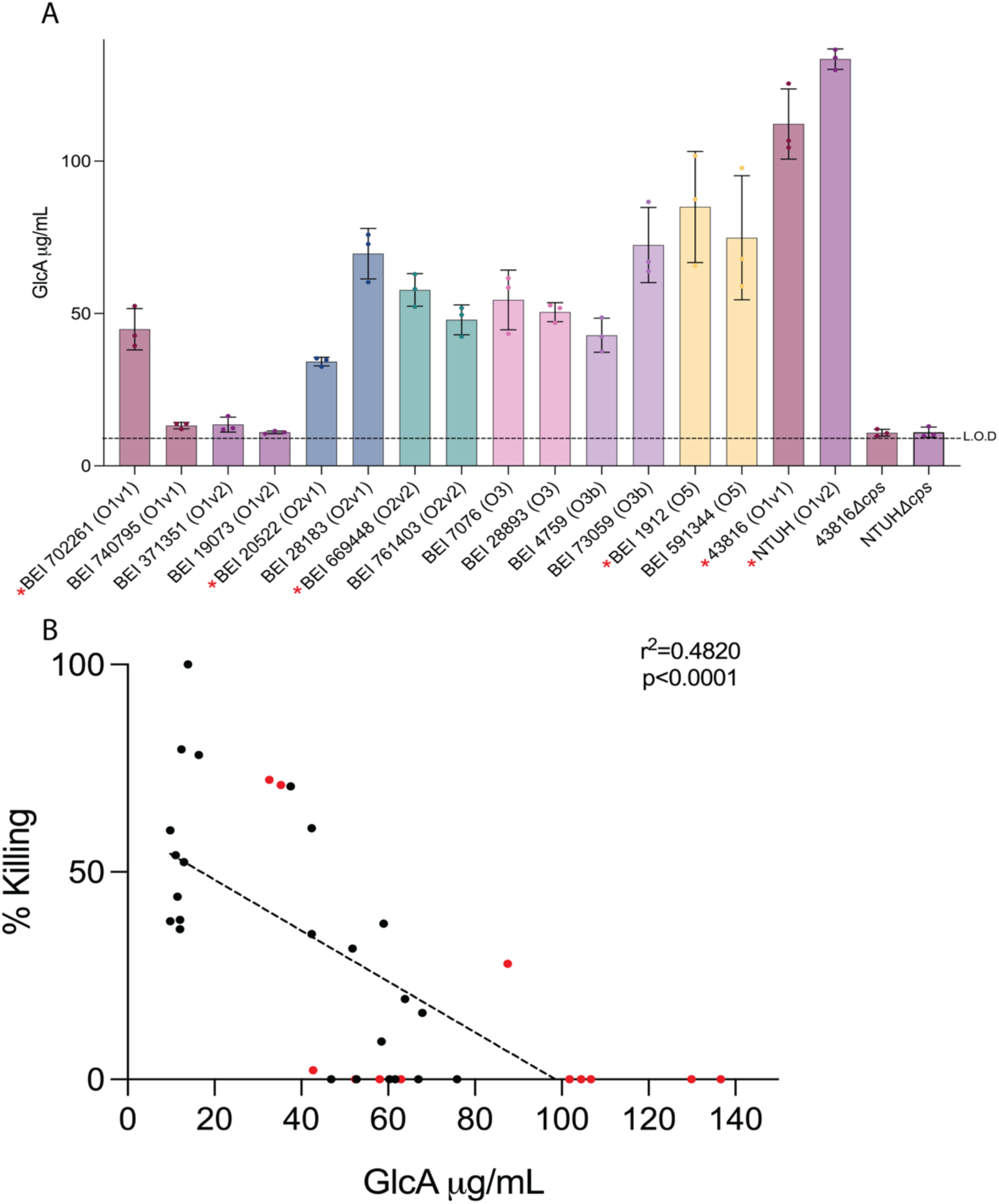
*K. pneumoniae* capsule quantification and correlation with serum bactericidality. (A) Capsule quantification using a glucuronic acid assay comparing 18 different strains of *K. pneumoniae*. The dotted line represents the limit of detection for uronic acids. (B) Simple linear regression analysis comparing the amount of capsule and the percent killing from serum bactericidal assays. Strains with the presence of hypervirulent genes are denoted with a red asterisk in A and red points in B. Non-linear regression was performed for B. Error bars represent means with standard deviations.

## Discussion

*K. pneumoniae* is an adaptable pathogen capable of causing an array of diseases in different hosts. More alarming, antibiotic resistance is on the rise, making this bacterium increasingly difficult to treat. *K. pneumoniae* is a common cause of nosocomial pneumonia (30) and is the most common cause of fatal sepsis in children under the age of five (4, 5). Vaccination represents an ideal preventive strategy for *K. pneumoniae* infections and would aid in lowering antibiotic use, potentially slowing the spread of antibiotic resistance. Currently, there is no licensed vaccine for *K. pneumoniae* on the market; however, one quadrivalent vaccine (Kleb4V) containing subtypes O1v1, O2v1, O2v2, and O3b is currently in human clinical trials (31). Here we present data on the production and functionality of a heptavalent O-antigen vaccine that includes most clinically relevant O-antigen subtypes and therefore represents the broadest polysaccharide vaccine targeting *K. pneumoniae* reported to date.

The last several years have demonstrated an increase in enthusiasm for *K. pneumoniae* vaccines with dozens of published preclinical studies, many with encouraging results. Several reports have focused on pure capsular polysaccharide vaccines or polysaccharide-based conjugate vaccines (22, 32–36). In addition to capsule, the components of LPS have also been considered for vaccine inclusion. Conjugate vaccines linking segments of LPS (other than the O-antigen) to a protein have been described (37, 38). These vaccines elicited antibodies; however, only one group demonstrated protection from a lethal *K. pneumoniae* challenge. Further, others have produced O-antigen conjugate vaccines using traditional chemical conjugation methods and demonstrated protection in murine models (39–42). Finally, using newer technologies others have created nanovaccines targeting the O1 and O2 O-antigens of *K. pneumoniae* (43, 44); however, the subtypes within these two serogroups were not investigated. In further support of O-antigen as a potential antigen target to combat *K. pneumoniae* infections, studies have documented protective monoclonal antibodies specific to O-antigen (45–49).

Here, we used the PglS bioconjugation technology to produce O-antigen conjugate vaccines against the broadest range of clinical isolates to date. Bioconjugation methods have certain advantages over traditional conjugation methods, particularly, it enables faster and easier production and therefore may be more cost effective. Further, it is easily adaptable for new polysaccharides or carrier proteins given its plasmid expression system within a glycoengineered bacterium. This last point is especially important as *K. pneumoniae* has numerous circulating strains and has the potential for serotype replacement following introduction of a vaccine into widespread use. The novel O-antigen bioconjugate vaccine reported here is the most comprehensive O-antigen vaccine targeting *K. pneumoniae* reported to date, containing three additional prevalent serotypes not included in other constructs.

Despite the functional and protective effects of O-antigen antibodies and vaccines reported in some studies, further work has demonstrated that O-antigen polysaccharide and O-antigen antibody binding can be blocked by the capsular polysaccharide in some strains (21, 28, 29). Indeed, here we find that higher capsule-producing strains exhibited diminished antibody binding and function compared to lower capsule-producing strains. Further, when capsule genes were deleted from such heavily encapsulated strains, binding and functionality of O-antibodies significantly increased. Our findings suggest that O-antigen antibodies may not successfully target all strains of *K. pneumoniae*, especially heavily encapsulated or hypervirulent strains.

In this current study we examined the functionality of a heptavalent *K. pneumoniae* O-antigen bioconjugate vaccine. To our knowledge this is the first conjugate vaccine to target not only the main serogroups of O-antigen, but also the prevalent subtypes within the groups as well. Our monovalent immunization studies suggest that each of the seven O-antigen subtypes we examined have distinct coverage and specificity that would partially be lost if they were omitted from a multivalent vaccine. New O-antigen subtypes continue to be discovered (14, 16) and must be tested for immunogenicity and cross-reactivity to determine appropriate inclusion in a polyvalent vaccine formulation.

We tested our vaccine-derived antibodies for binding and function across a diverse panel of 18 different strains of *K. pneumoniae* that express one of the seven O-antigen subtypes, in addition to a variety of different capsule types at varying amounts. While vaccine-derived antibodies were able to bind and induce complement-mediated killing in roughly half of the strains, the other strains demonstrated little to no binding or functionality. We hypothesize this is due to increased capsule quantity and subsequent antibody blockage by the capsule as indicated by a negative correlation between these two measurements. However, we also provide evidence in this study for clinical strains that produce little capsule; thus, capsule-targeting antibodies may also yield poor results in these isolates. Therefore, in the context of a vaccine, we hypothesize that an ideal vaccine against *K. pneumoniae* may include several of the described O-antigen types in addition to some of the most prevalent capsular types. Future studies should consider multivalent O-antigen and capsule vaccines. Further, any vaccine candidate will require testing against numerous *K. pneumoniae* strains expressing a variety of different O-antigen and capsule types, in addition to differing amounts of capsule. These studies are imperative to developing an efficacious multivalent vaccine to target this increasingly resistant pathogen.

## Materials and Methods

### Bacterial strains, plasmids, and growth conditions

*K. pneumoniae* strains used in this work are listed in Table 1. *K. pneumoniae* strains and mutants were used for all ELISAs and serum bactericidal assays. Fourteen of the 18 strains used for this study are part of the the BEI Resources Repository *Klebsiella pneumoniae* diversity panel (27) which was obtained from BEI Resources, NIAID, NIH: *Klebsiella pneumoniae* MRSN Diversity Panel, NR-55604, provided by the Multidrug-Resistant Organism Repository and Surveillance Network (MRSN) at the Walter Reed Army Institute of Research (WRAIR), Silver Springs, Maryland, USA. BEI strains were categorized according to the BEI website for their O-antigen and capsule. ATCC 43816 was previously determined as capsule type K2 and O1v1 O-antigen type (50). NTUH-K2044 was previously determined as capsule type K1 and O1v2 O-antigen type (51). Other bacterial strains used for the expression of the bioconjugates are listed in Supplemental Table 1.

### Construction of O-antigen polysaccharide expression plasmids

DNA primers used for PCR are listed in Supplemental Table 2. All PCR products were gel-purified prior to Gibson assembly into expression vectors. The *wzm* – *wbbO* genes used to produce O-antigens O2v1, O2v2, and O1v1 were amplified from purified NTUH gDNA (Wizard Genomic DNA Purification Kit, Promega) using primers O2a_cluster_1F/O2a_cluster_1R. The *wzm* – *wbbO* PCR product was assembled with PCR-linearized pBBR1MCS2 plasmid (Addgene, primers pBBR1-1F/pBBR1-1R) using an NEBuilder HiFi DNA Assembly kit (NEB). The *gmlABC* genes used to make serotype O2v2 were PCR-amplified from NTUH gDNA using primers gmlABC_1F/gmlABC_1R and assembled with XbaI restriction enzyme-linearized pACT3 vector. The *wbbY* gene used to make serotype O1v1 was PCR-amplified from NTUH gDNA using primers wbbY_1F/wbbY_1R and assembled with XbaI restriction enzyme-linearized pACT3 vector. The O3 gene cluster was PCR-amplified from *E. coli* O9:K9:H12 (SSI Diagnostica, Strain 81842) in two fragments using primer pairs O3_1F/O3_1R and O3_2F/O3_2R. The O3b gene cluster was PCR-amplified in two fragments from *K. pneumoniae* TOP52 using primer pairs O3_1F/O3b_1R and O3b_2F/ O3_2R. The O5 gene cluster was PCR-amplified from *E. coli* O8:K8:H4 (SSI Diagnostica, Strain 81841) in two fragments using primer pairs O3_1F/O5_1R and O5_2F/O3_2R. Each pair of DNA fragments composing the O3, O3b or O5 gene clusters were Gibson assembled with PCR-linearized pBBR1MCS2 plasmid (primers pBBR1-1F/pBBR1-1R). After assembly, the Gibson assembly reactions were transformed into chemically competent *E. coli* Stellar cells (Takara Bio) and plated on LB agar supplemented with antibiotics. All plasmids were sequence-verified by Sanger sequencing (Genewiz) prior to use.

### Production of bioconjugates

O2v1 O-antigen was produced in *E. coli* by expressing pBBR1-MCS2-*wzm* – *wbbO*. O2v2 was produced by co-expression of pBBR1-MCS2-*wzm* – *wbbO* and pACT3-*gmlABC*. O1v1 was produced by co-expression of pBBR1-MCS2-*wzm* – *wbbO* and pACT3-*wbbY*. O3, O3b, and O5 were produced by expression of pBBR1MCS2-O3, pBBR1MCS2-O3b, or pBBR1MCS2-O5, respectively. Sequence-verified O-antigen plasmids were transformed into *E. coli* CLM24 (26) carrying plasmid pVNM245 that encodes *Pseudomonas aeruginosa* EPA with two internal glycotags and downstream *Acinetobacter baylyi* ADP1 PglS OTase (21, 25). pVNM245 and PglS OTase plasmids were transformed into NTUH1*wcaJ*1*waaL* for expression of O1v2. Overnight transformants were selected on LB agar media supplemented with ampicillin (100 μg/mL), kanamycin (20 μg/mL) and/or broth media. The next day, starters were diluted into fresh TB media (1 L media in 2 L non-baffled flasks) to a starting OD_600_ of 0.05. All strains were cultured at 30 °C to a mid-log OD_600_ of 0.3 – 0.5 before induction with 0.1 mM IPTG and overnight culturing. All bioconjugates were produced at 30°C post-induction except O1v1 and O2v2, which were produced at 25 °C. The next morning, cells were pelleted by centrifugation at 7,000 *xg* and stored at –20 °C until purification. Unglycosylated EPA control protein was expressed from pVNM245 in the absence of O-antigen plasmids.

### Purification of EPA and O-antigen bioconjugates and characterization

Each of the bioconjugates were purified from *E. coli* periplasmic extracts using previously described methods (21). Protein purity was assessed using western blotting and Coomassie-stained protein gels as previously described (21). Western blots were probed with mouse O-antigen antisera except for O1v1, which was detected by rabbit antisera (antisera were gifts from Prof. Chris Whitfield). The protein antibody used for western blotting was mouse or rabbit anti-6xHis antibodies (Millipore-Sigma). Secondary antibodies were IRDye 680RD goat anti-mouse and/or IRDye 800CW goat anti-rabbit (Li-Cor). Protein concentrations of the purified bioconjugates were determined using BCA Protein Assay kit (Pierce) and their sugar content quantified using an anthrone-sulfuric acid assay.

### Nuclear magnetic resonance (NMR) spectroscopy

Polysaccharides heterologously expressed in E. coli were extracted by heating whole-cell bacteria in 2% acetic acid at 105°C for 1.5 hours. Insoluble material was removed by centrifugation, and the supernatant, containing a polysaccharide hydrolyzed from the core saccharide of E. coli, was subjected to purification on a Sephadex G50 column, subsequently purified on Hitrap Q (acidic) column, and polished on a Hitrap S column to remove trace protein background. Polysaccharides were dried and analyzed by NMR. NMR experiments were carried out on a Bruker AVANCE III 600 MHz ( 1 H) spectrometer with 5 mm Z-gradient probe with acetone internal reference (2.23 ppm for 1 H and 31.45 ppm for 13C) using standard pulse sequences cosygpprqf (gCOSY), mlevphpr (TOCSY, mixing time 120 ms), roesyphpr (ROESY, mixing time 500 ms), hsqcedetgp (HSQC), hsqcetgpml (HSQC-TOCSY, 80 ms TOCSY delay) and hmbcgplpndqf (HMBC, 100 ms long range transfer delay). Resolution was kept <3 Hz/pt in F2 in proton-proton correlations and<5 Hz/pt in F2 of H-C correlations. The spectra were processed and analyzed using the Bruker Topspin 2.1 program. Monosaccharides were identified by COSY, TOCSY, and NOESY cross peak patterns and 13C NMR chemical shifts. Connections between monosaccharides were determined from transglycosidic NOE and HMBC correlations.

### Mass spectrometry

Intact mass analysis was performed on a 6520 Accurate Mass Quadrupole Time-of-Flight Mass Spectrometer coupled to an Agilent 1200 Infinity HPLC (Agilent, Santa Clara, CA, USA) using a Jupiter 300 C5 column (2mm*50mm, Phenomenex). Protein samples were resuspended in 2% acetonitrile, 0.1% trifluoroacetic acid and loaded/separated on the C5 column at a flow rate of 0.25 mL/min. 2 to 5 μg of each bioconjugate was desalted on column for 2 min with Buffer A (2% acetonitrile 0.1% formic acid) before being separated by altering the percentage of Buffer B (80% acetonitrile, 0.1% formic acid) from 0% to 100% over 16.5 min. The column was then held at 100% Buffer B for 0.5 min before being equilibrated for 1 min with Buffer A, for a total run time of 20.0 min. Samples were infused into the Time-of-Flight Mass Spectrometer using electrospray ionization (ESI) and MS1 mass spectra acquired with a mass range of 300–3000m/z at 1 Hz. Intact mass analysis and deconvolution was performed using MassHunter B.06.00 (Agilent).

### Construction of K. pneumoniae mutant strains

A modified Red recombinase protocol was utilized to construct the Δ*cps* mutants (lacking the *cps* operon from *wzi* to *wcaJ*) using pKD46s (21, 52). Linear regions of DNA were amplified from pKD4 using primers listed in Supplemental Table 2, resulting in the region of interest being replaced with a kanamycin cassette. The kanamycin cassette was then removed using pCP20. All mutants were confirmed by sequencing of amplicons generated via PCR using specific check primers. The *wcaJ* and *waaL* mutant was made using a modified CRISPR protocol (53). The zeocin resistance cassette of pUC19_CRISPR_DpmrA was replaced with a tetracycline resistance gene, and the *pmrA* gRNA and homology arms were swapped for those targeting *wcaJ* or *waaL*. Deletion mutants were identified by PCR and confirmed by sequencing.

### Murine vaccination

All murine immunizations complied with ethical regulations for animal testing and research. Experiments were carried out at Washington University School of Medicine in St. Louis according to the institutional guidelines and received approval from the Institutional Animal Care and Use Committee at Washington University in St. Louis. Five-week-old male and/or female CD-1 mice (Charles River) were subcutaneously injected with 100 μL of a vaccine formulation on days 0, 14, and 28. The vaccination groups were as follows: EPA carrier protein alone, O1v2-EPA, O1v2-EPA, O2v1-EPA, O2v2-EPA, O3-EPA, O3b-EPA, O5-EPA, and O7V-EPA, which is a heptavalent mixture of all O-antigen bioconjugates. All vaccines were formulated with Aldydrogel® 2% aluminum hydroxide gel (InvivoGen) at a 1:9 ratio (50 μL vaccine to 5.5 μL adjuvant in 44.5 μL sterile PBS). All vaccination groups received 1 μg of vaccine based on total polysaccharide content. The total polysaccharide content was measured using a modified anthrone-sulfuric assay. Sera were collected on days 0, 14, 28, and 42 prior to immunizations.

### Enzyme-linked immunosorbent assays

Briefly, 96 well plates (BRAND^TM^ immunoGrade microplates) were coated overnight with ∼10^6^ CFU/100 μL of the specified *K. pneumoniae* strain in sodium carbonate buffer. All strains and growth conditions are described above. After coating, wells were blocked with 1% BSA in sterile PBS and washed with 0.05% PBS-Tween-20 (PBS-T); all subsequent washes were the same. Sera from immunized mice were diluted 1:100 and added to wells in triplicate or quadruplicate for 1 h at room temperature. After washing, HRP-conjugated anti-mouse IgG (GE Lifesciences, 1:5000 dilution in PBS-T) was added to wells for 1 h at room temperature. Plates were washed, developed using 3,3’,5,5’-tetramethyl benzidine (TMB) substrate (Biolegend), and stopped with 2 N H_2_SO_4_. Absorbance was determined at 450 nm using a microplate reader (Agilent). Total IgG concentration was determined using an IgG standard curve. Standard wells were coated with diluted mouse IgG in sodium carbonate buffer and treated the same as sample wells thereafter. All wells were normalized to blank wells coated and treated the same as sample wells without receiving primary mouse sera.

### Serum bactericidal assays

Serum bactericidal assays for *K. pneumoniae* were adapted from methods previously described (21). *K. pneumoniae* isolates and derived mutants were grown under static conditions in LB broth for 16 hours at 37°C. Cultures were centrifuged at 8000 *x g* for 10 min and resulting pellets were resuspended in sterile PBS to an OD_600_ ∼1.0. Cultures were diluted 1:10,000 in sterile PBS. The assay mixture was prepared in a 96-well U-bottom microtiter plate (TPP®) by combining 70 μL of diluted bacteria and 20 μL of diluted heat-inactivated mouse serum in triplicate per bacterial strain. Pooled day 42 sera from mice immunized O7V-EPA bioconjugate were heat inactivated at 56 °C for 30 min. After incubation at 37 °C with shaking for 1 h, 10 μL of baby rabbit complement (Pel-Freez Biologicals) was added to wells at a final concentration of 10% and incubated for an additional 1 h at 37 °C with shaking. Control wells were treated the same as samples except for receiving diluted, heat-inactivated, pre-immune mouse serum. After the final incubation, samples were serially titrated in sterile PBS and plated in pentaplicate. Colonies were counted after 16-h incubation at room temperature. SBA titers were reported as percent killing compared to control.

### Glucuronic acid quantification

Quantification of capsule was performed using glucuronic acid assays. Briefly, bacteria were grown overnight in LB broth at 37 °C under static growth conditions. Cultures were pelleted and resuspended in 1x PBS to an OD_600_ of 1.5. A total of 500 μL of normalized culture was mixed with 100 μL of 1% Zwittergent 3-14 (Sigma) in triplicate in 100 mM citric acid and incubated at 50 °C for 20 min. After centrifugation, supernatants from samples were precipitated with cold ethanol at 4 °C for 20 min. Upon precipitation, samples were recentrifuged and the pellets were dissolved in 200 μL sterile water and 1200 μL of 12.5 mM tetraborate in concentrated H_2_SO_4_. Samples were vortexed, boiled at 95 °C for 5 min, and mixed with 20 μL of 0.15% 3-hydroxydiphenol (Sigma) in 0.5% NaOH. Absorbance was measured at 520 nm using a microplate reader (Agilent). The uronic acid concentration of each sample was determined using a standard curve of glucuronic acid (Sigma). The limit of detection (LOD) was previously defined (54).

### Statistics

All statistics were performed in Graphpad Prism 10. Mann-Whitney U tests were used to analyze ELISA, killing and capsule quantification significance, as data were not all normally distributed. Simple linear regression analysis was used for comparison of capsule quantities and percent bacterial killing by serum bactericidal assays. All errors bars are shown as means with standard deviations. p<0.05 was considered significant.

## Acknowledgements

We would like to thank Dr. Chris Whitfield (University of Guelph) for the generous gift of *K. pneumoniae* O-antigen animal antisera and Dr. David Hunstad (Washington University) for criticial review of this manuscript.

