## Supplemental Figures for "A heptavalent O-antigen bioconjugate vaccine exhibits differential functional antibody responses against diverse *Klebsiella pneumoniae* isolates"

### Supplemental Figures and Tables

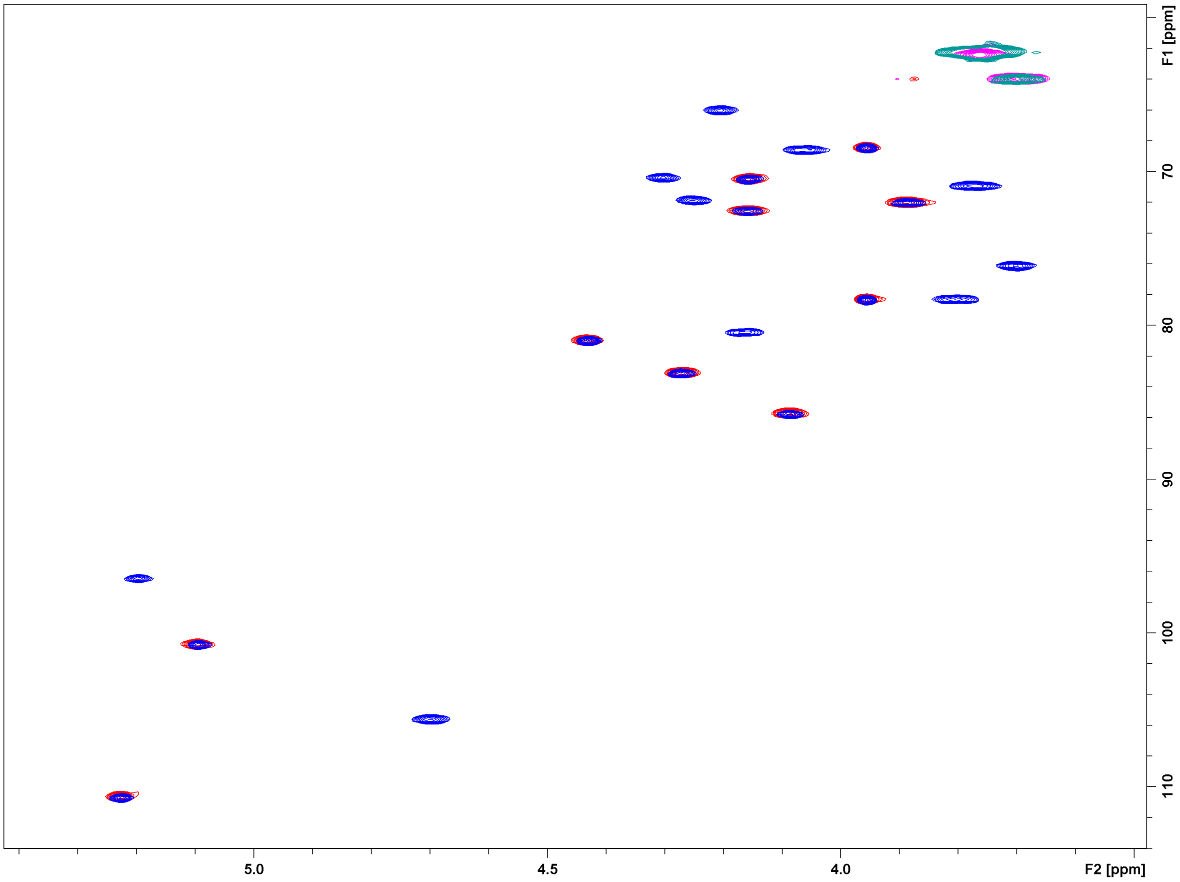


**Supplemental Figure 1.** NMR of *K. pneumoniae* O2a (O2v1) O-antigen polysaccharide linked to *E. coli* core saccharide. Spectral comparison of the $^{1}H$-$^{13}C$ HSQC spectra of native *K. pneumoniae* O1 polysaccharide (blue-cyan) overlayed with the O2a polysaccharide produced by glycoengineered *E. coli* (red-pink).

A

```
 A          B
-3-β-Galf-3-α-Gal-
            |4
         α-Gal
           C
```

| Unit | | H/C-1 | H/C-2 | H/C-3 | H/C-4 | H/C-5 | H/C-6a;b |
|---|---|---|---|---|---|---|---|
| β-Galf A | H | 5.23 | 4.35 | 4.09 | 4.31 | 3.87 | 3.70; 3.70 |
| | C | 110.8 | 81.6 | 85.9 | 80.9 | 71.1 | 64.0 |
| α-Gal B | H | 5.10 | 4.10 | 3.96 | 4.19 | 4.18 | 3.85; 3.92 |
| | C | 101.2 | 68.9 | 77.8 | 79.4 | 73.4 | 61.4 |
| α-Gal C | H | 5.02 | 3.84 | 3.92 | 4.08 | 4.23 | 3.80; 3.82 |
| | C | 101.4 | 70.0 | 70.2 | 69.8 | 71.8 | 61.2 |

B

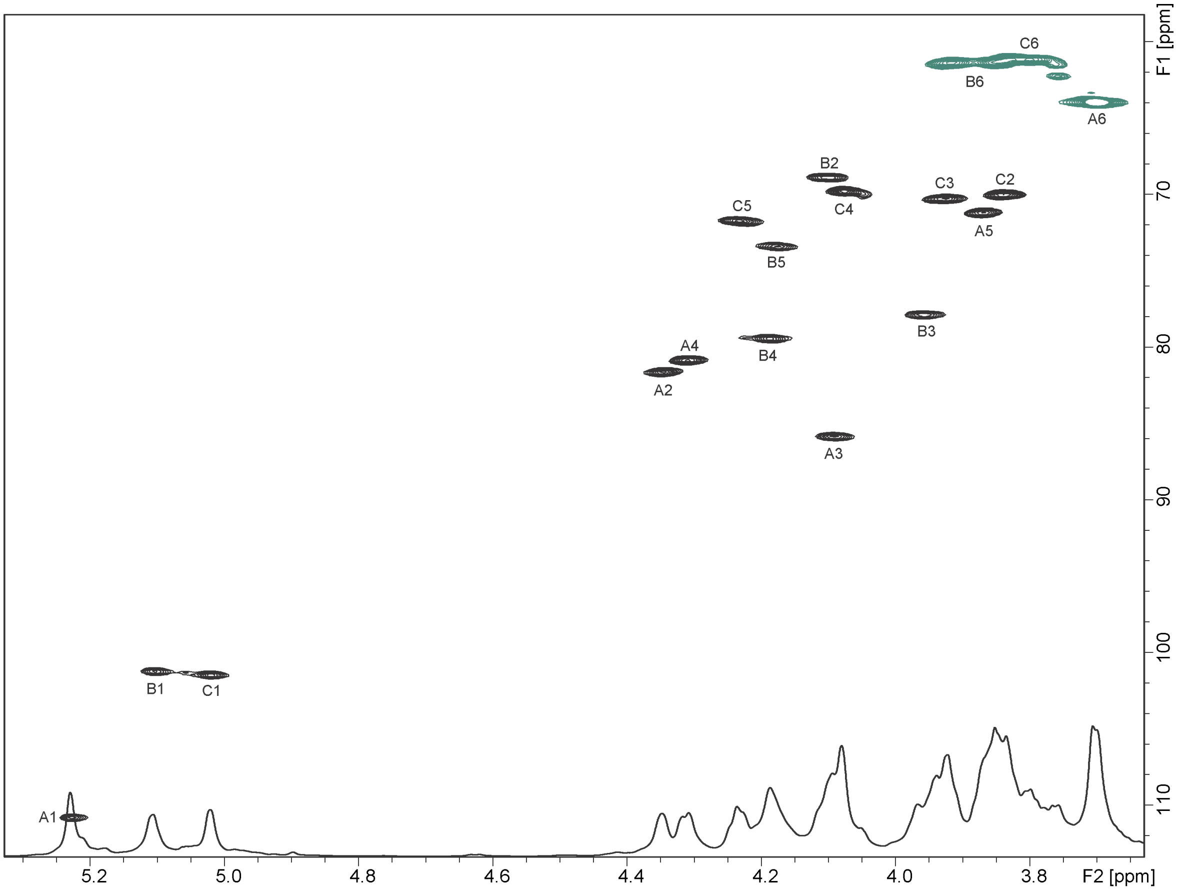


**Supplemental Figure 2.** NMR of *K. pneumoniae* O2afg (O2v2) polysaccharide linked to *E. coli* core saccharide. NMR data (A) and spectral comparison (B) of the $^{1}$H-$^{13}$C HSQC spectra of native *K. pneumoniae* O2afg was identical to the known Galactan-III structure.

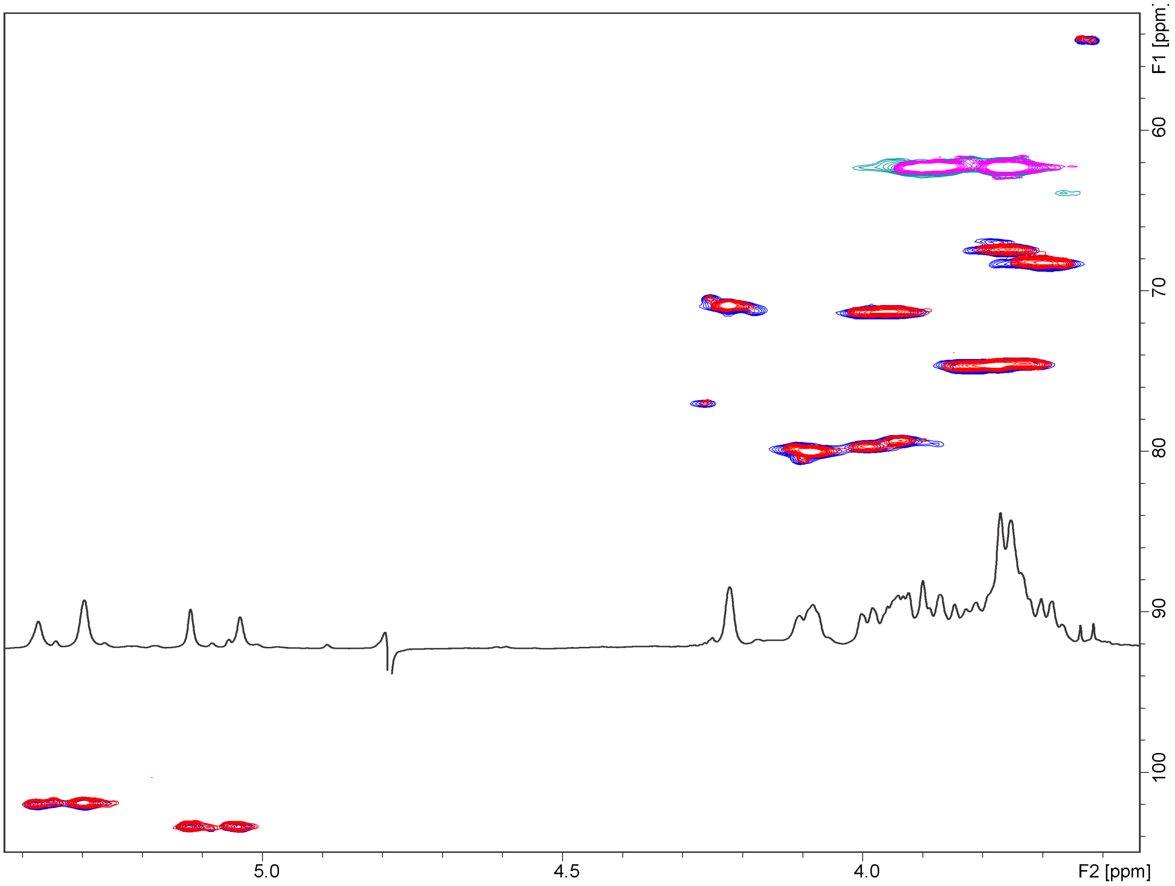


**Supplemental Figure 3.** NMR of *K. pneumoniae* O3 O-antigen polysaccharide linked to *E. coli* core saccharide. Spectral comparison of the $^{1}H$-$^{13}C$ HSQC spectra of native *K. pneumoniae* O3 polysaccharide (blue-cyan) was identical to the polysaccharide produced by glycoengineered *E. coli* (red-pink), including the terminal methylphosphate-3-mannose cap.

A Main chain repeating structure:

-2-α-Man-3-α-Man-3-α-Man-
A B C

Non-reducing end terminal modification:

Me*P*-3-α-Man-2-α-Man-3-α-Man-3-α-Man-
E D B C

| Unit | | H/C-1 | H/C-2 | H/C-3 | H/C-4 | H/C-5 | H/C-6a;b |
|---|---|---|---|---|---|---|---|
| α-Man A | H | 5.38 | 4.11 | 4.00 | 3.69 | 3.79 | 3.76-3.91 |
| | C | 101.9 | 79.5 | 71.2 | 68.2 | 74.6 | 62.1 |
| α-Man B | H | 5.13 | 4.23 | 4.01 | 3.78 | 3.79 | 3.76-3.91 |
| | C | 103.3 | 70.8 | 79.4 | 67.3 | 74.6 | 62.1 |
| α-Man C | H | 5.05 | 4.23 | 3.95 | 3.78 | 3.79 | 3.76-3.91 |
| | C | 103.3 | 70.8 | 79.2 | 67.3 | 74.6 | 62.1 |
| α-Man D | H | 5.43 | 4.10 | 3.97 | 3.78 | 3.78 | 3.76-3.91 |
| | C | 101.7 | 80.3 | 71.2 | 68.2 | 74.6 | 62.1 |
| α-Man E | H | 5.06 | 4.26 | 4.28 | 3.81 | 3.85 | 3.76-3.91 |
| | C | 103.3 | 70.4 | 76.9 | 66.7 | 74.6 | 62.1 |

B

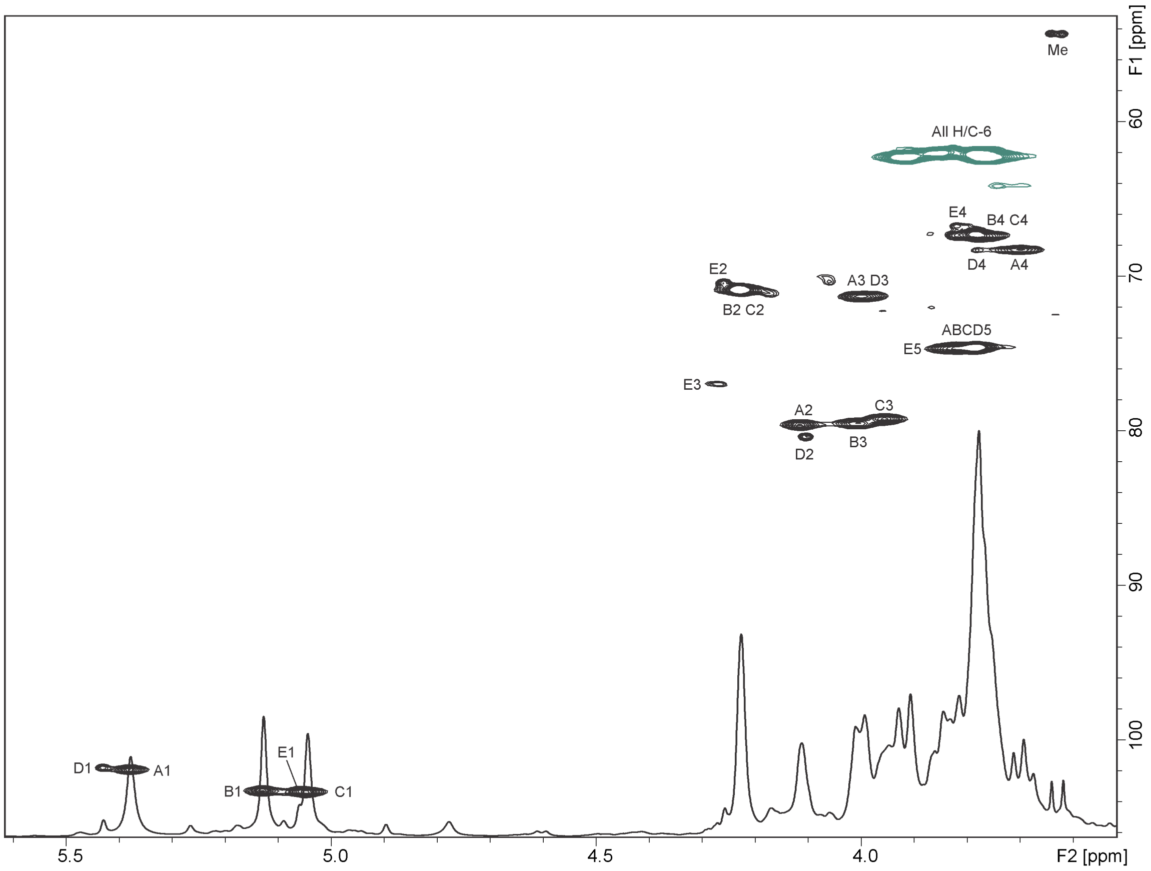


**Supplemental Figure 4.** NMR of *K. pneumoniae* O3b polysaccharide linked to *E. coli* core saccharide. NMR data (A) and $^{1}$H-$^{13}$C HSQC spectrum (B) of *K. pneumoniae* O3b polysaccharide with P position confirmed by $^{1}$H-$^{31}$P HSQC and $^{1}$H-$^{31}$P HMQC-TOCSY.

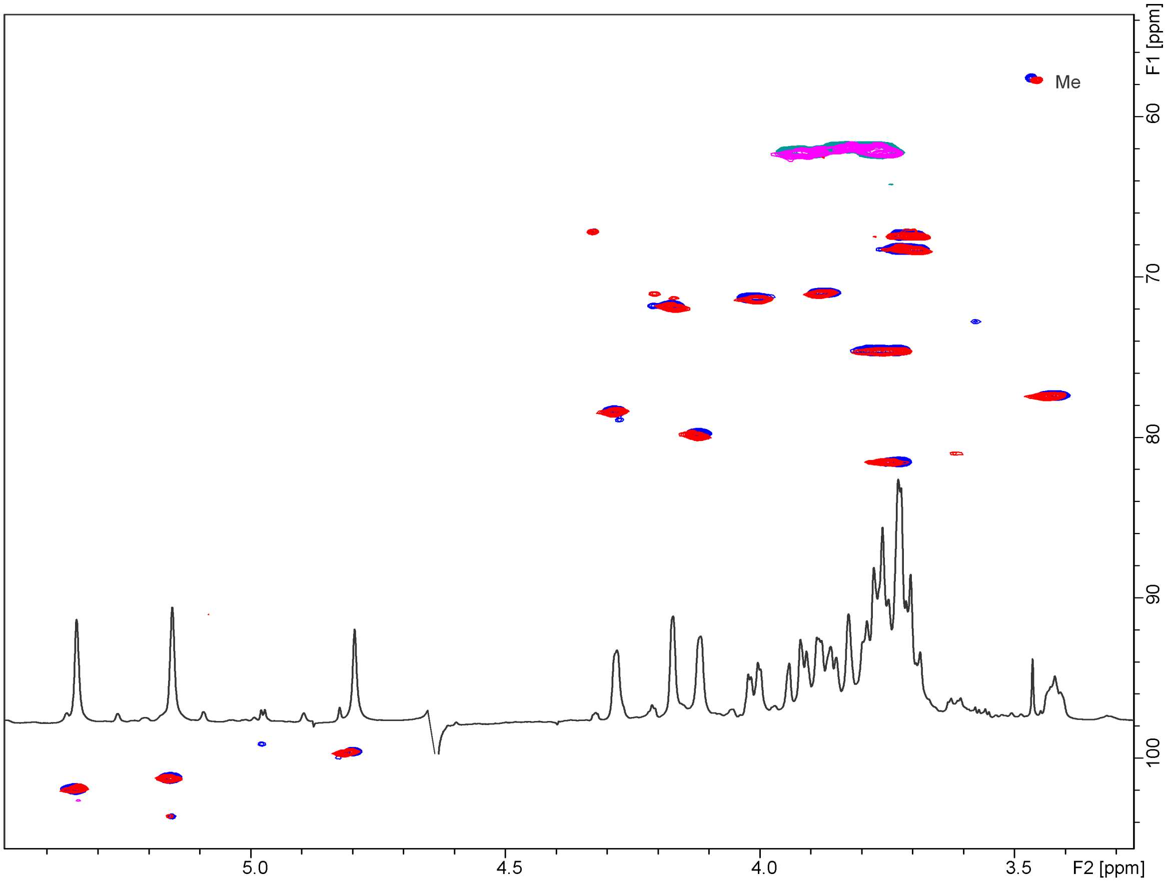


**Supplemental Figure 5.** NMR of *K. pneumoniae* O5 O-antigen polysaccharide linked to *E. coli* core saccharide. Spectral overlap of the $^{1}H$-$^{13}C$ HSQC spectra of native *K. pneumoniae* O5 polysaccharide (blue-cyan) was identical to the polysaccharide produced by glycoengineered *E. coli* (red-pink), including the terminal methyl cap.

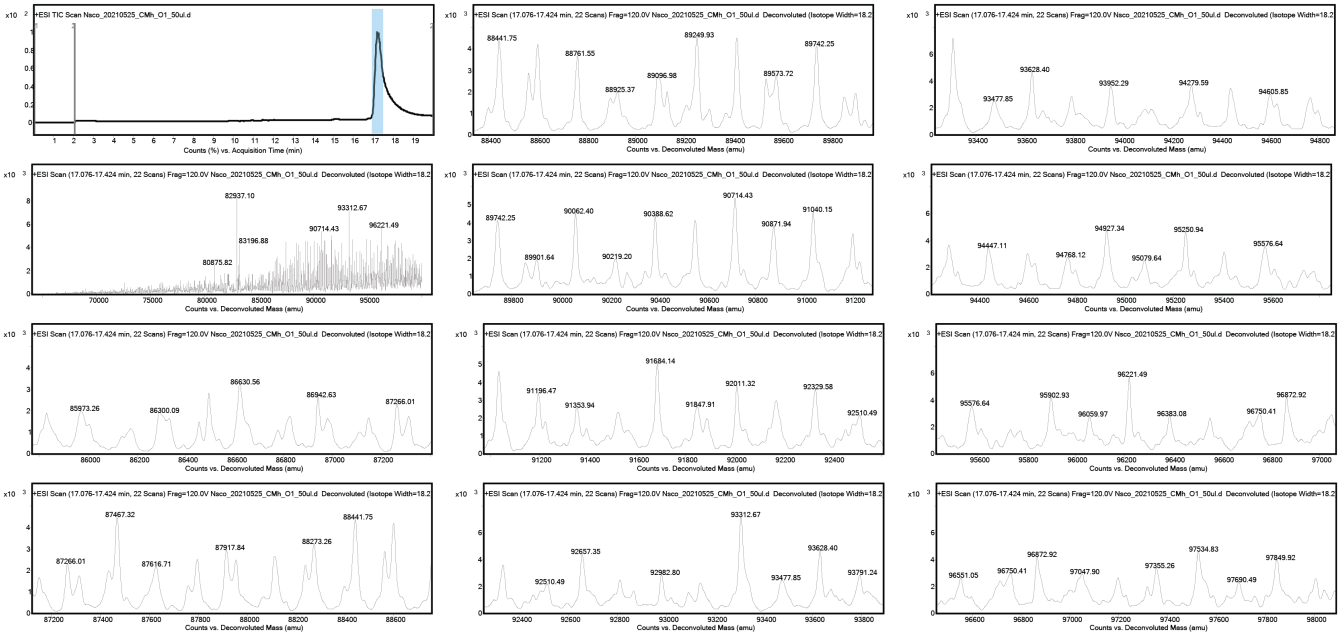

**Supplemental Figure 6.** MS1 spectrum of intact, purified O1v1-EPA.

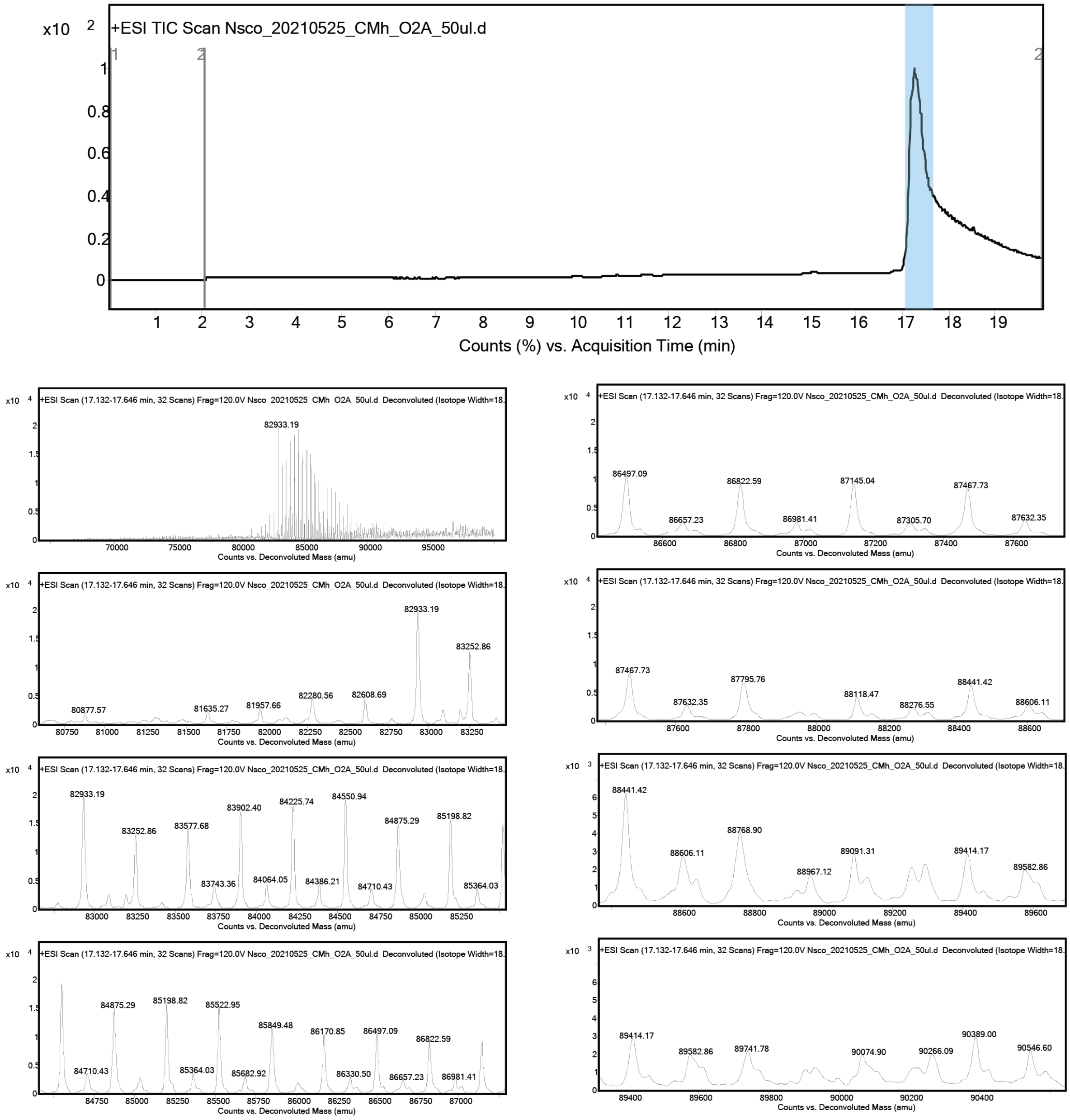


**Supplemental Figure 7.** MS1 spectrum of intact, purified O2v1-EPA.

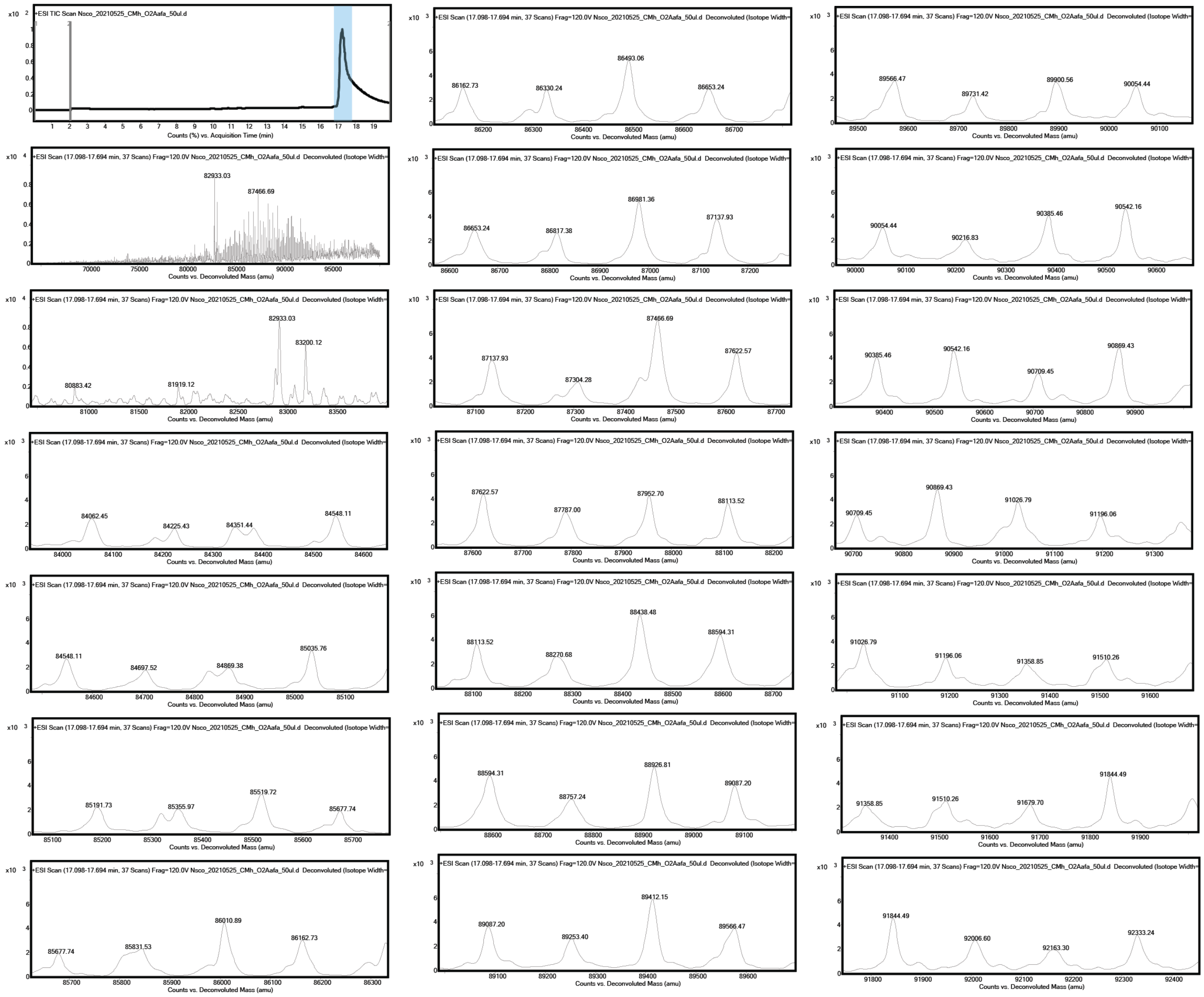


**Supplemental Figure 8.** MS1 spectrum of intact, purified O2v2-EPA.

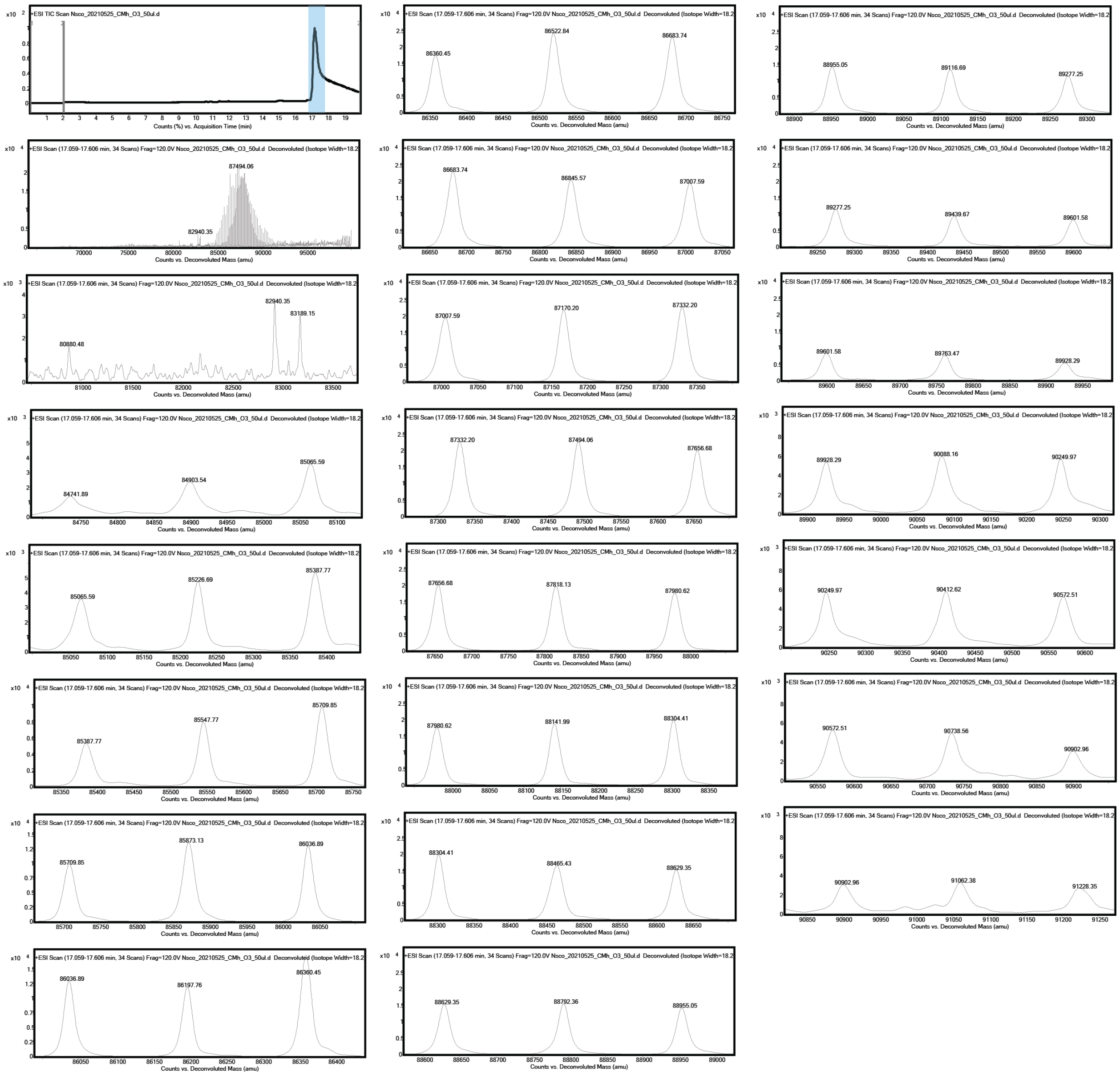

**Supplemental Figure 9.** MS1 spectrum of intact, purified O3-EPA.

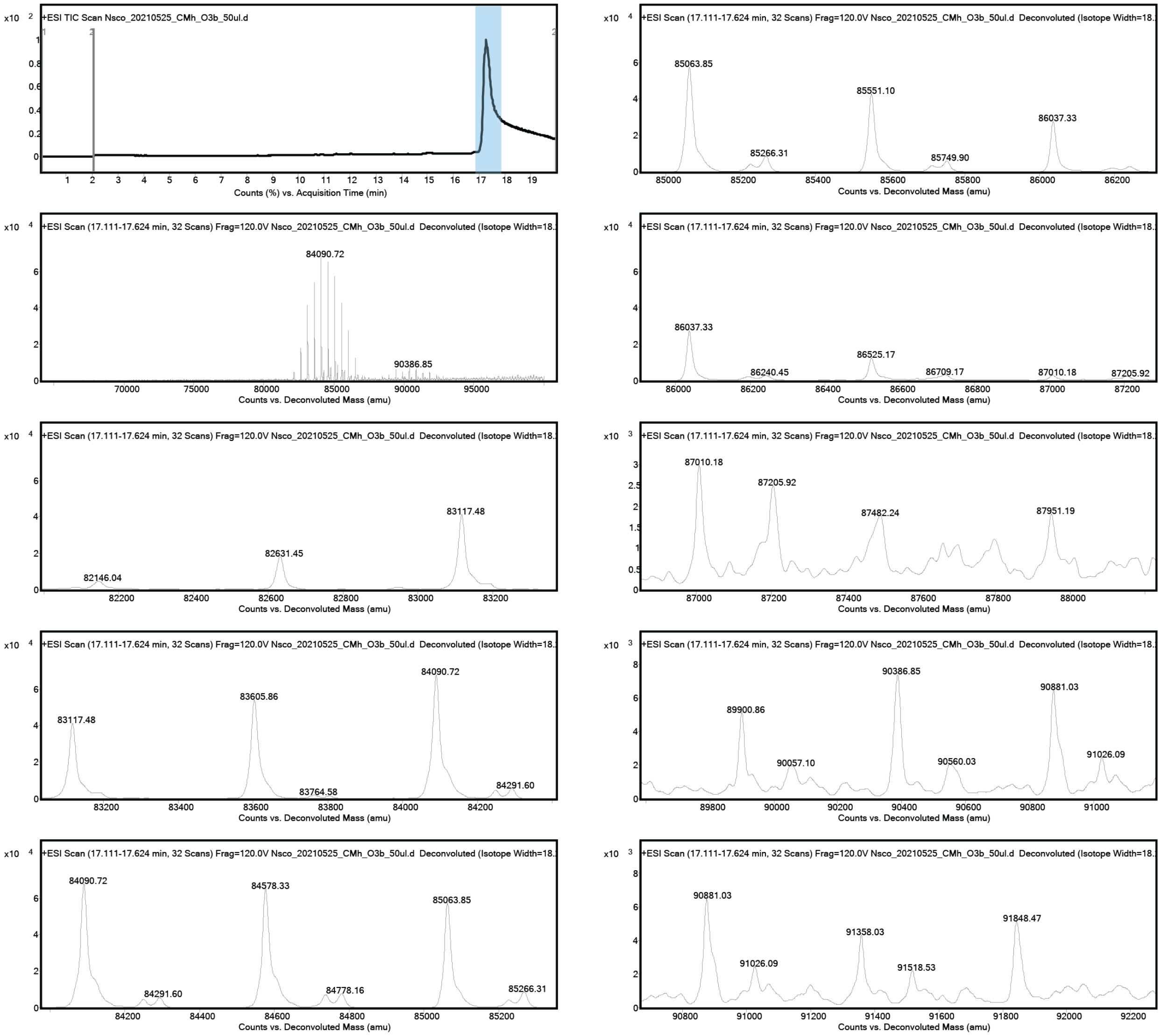


**Supplemental Figure 10.** MS1 spectrum of intact, purified O3b-EPA.

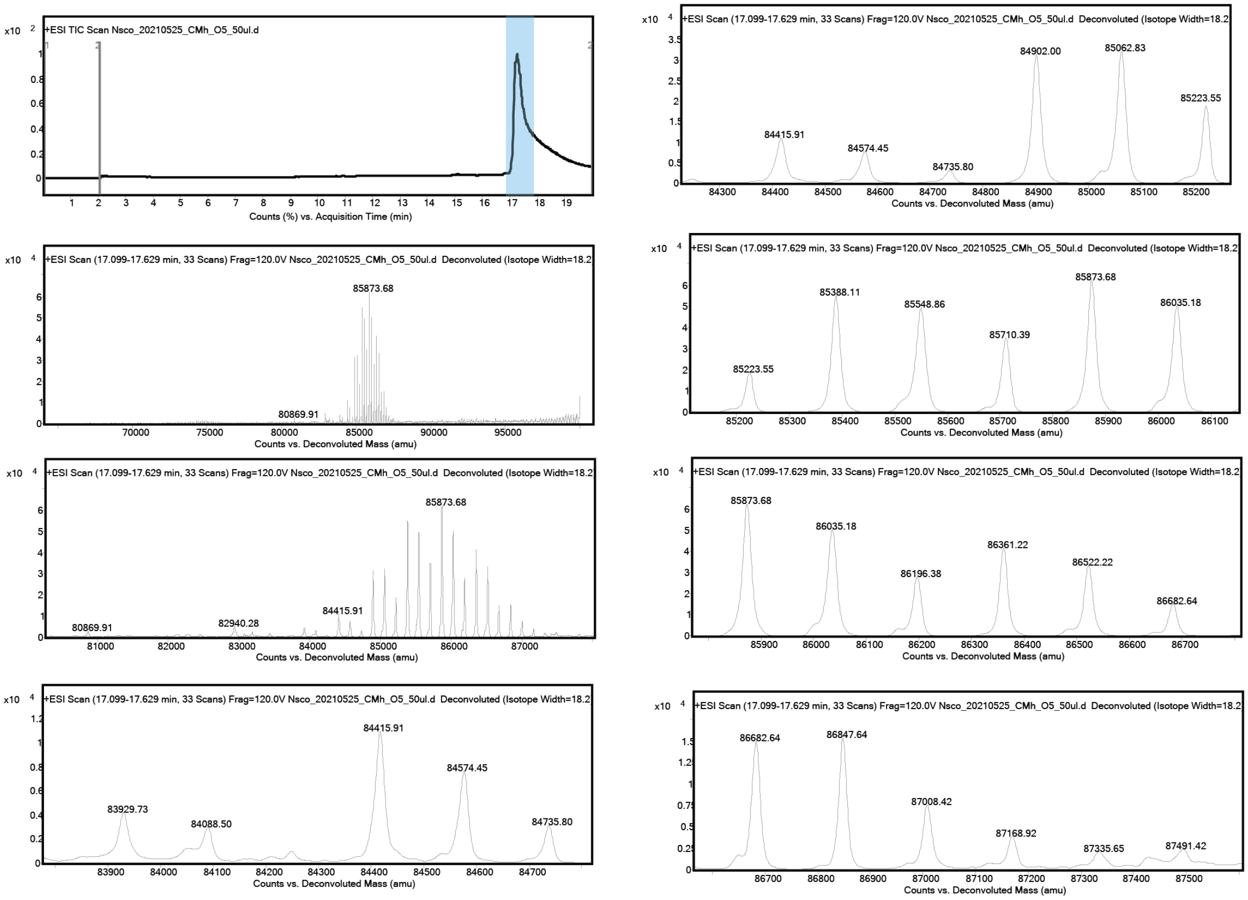


**Supplemental Figure 11.** MS1 spectrum of intact, purified O5-EPA.

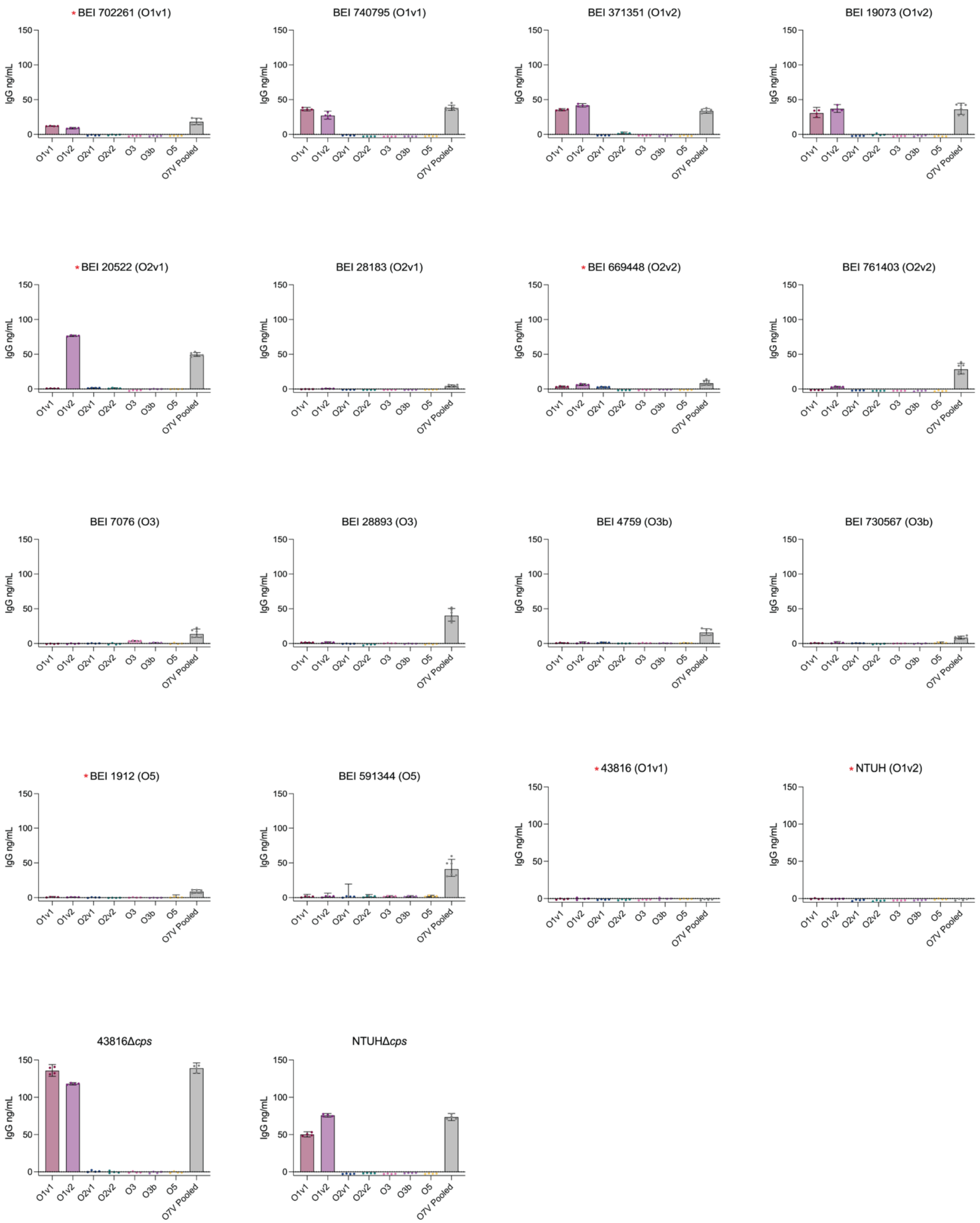


**Supplemental Figure 12.** Detection of IgG generated by immunization of CD-1 mice with monovalent or heptavalent O-antigen bioconjugates against strains of *K. pneumoniae*. O-antigen specific IgG from day 42 serum as measured by ELISA against 18 different strains of *K. pneumoniae*. Error bars represent means with standard deviations.

| Strain or Plasmid | Genotype/plasmid genes | Reference |
|---|---|---|
| E. coli CLM24 | W3110ΔwaaL | (25) |
| pVNM245 | *P. aeruginosa* EPA and A. baylyi ADP1 PglS expressed from pEXT20 backbone, $Amp^R$ | (20) |
| pVNM320 | *P. aeruginosa* EPA and A. baylyi ADP1 PglS expressed from pEXT20 backbone, $Tet^R$ | This work |
| pBBR1-MCS2-wzm – wbbO | NTUH O-antigen cluster (wzm, wzt, wbbM, glf, wbbN, wbbO) expressed from pBBR1-MCS-2 backbone, $Kan^R$ | This work, (20) |
| pACT3-gmlABC (pVNM39) | NTUH gmlABC expressed from pACT3 plasmid backbone, $Cml^R$ | This work |
| pACT3-wbbY (pVNM40) | NTUH wbbY expressed from pACT3 plasmid backbone, $Cml^R$ | This work, (20) |
| pBBR1-MCS2-O3 (pVNM76) | *E. coli* O9:K9:H12 O3 O-antigen cluster (manC – wbdC) expressed from pBBR1-MCS-2 backbone, $Kan^R$ | This work |
| pBBR1-MCS2-O3b (pVNM59) | *K. pneumoniae* TOP52 O3b O-antigen cluster (manC – wbdC) expressed from pBBR1-MCS-2 backbone, $Kan^R$ | This work |
| pBBR1-MCS2-O5 (pVNM71) | *E. coli* O8:K8:H4 O-antigen cluster (manC – wbdC) expressed from pBBR1-MCS-2 backbone, $Kan^R$ | This work |
| NTUHΔ*wcaJ*Δ*waaL* | *K. pneumoniae* strain used for production of O1v2 O-antigen | This work |
| TOP52 | *K. pneumoniae* strain used for cloning O3b | This work |

**Supplemental Table 1.** Additional bacterial strains and plasmids used in this study.

| Primer | Sequence |
|---|---|
| **PCR primers for cloning *K. pneumoniae* O-antigen genes and vectors** | |
| pBBR1-1F | gcgttaatattttgttaaaattcgcg |
| pBBR1-1R 101 | agctgtttcctgtgtgaaattg |
| O2a_cluster_1F | gcggataacaatttcacacaggaaacagctatgaagtacaatttagggtatttatttg |
| O2a_cluster_1R | ttaacgcgaattttaacaaaatattaacgctcatcgaactacatcatgatatatttg |
| gmlABC_1F | ctcggtacccggggatcctcatgccaagttcaggccc |
| gmlABC_1R | atgcctgcaggtcgactctactaataatttatcgttgaccttcgc |
| wbbY_1F | atgcctgcaggtcgactctattattttaacattgatttcactttccgg |
| wbbY_1R | ctcggtacccggggatcctcatgaagaaaattcttataatgacgcc |
| O3_1F | gcggataacaatttcacacaggaaacagctatgttacttcctgtaattatgg |
| O3_1R | accagcattttcaataattgc |
| O3_2F | atgagccgcgcaattattg |
| O3_2R | ttaacgcgaattttaacaaaatattaacgctcaggatttgctttccag |
| O3b_1R | gccaatcgaaccaaaatc |
| O3b_2F | gctcgtttaatagattttggttcg |
| O5_1R | aatacgcatcgttatttctc |
| O5_2F | taaaaatacggagaaataacgatg |
| **Primers used to generate *K. pneumoniae* capsule knockouts** | |
| 43816 wzi Fwd | aaaatggatcttgtacaatgataaaaattgcgcgcattgccgtgagtgtaggctggagctgcttc |
| 43816 wcaJ Rvs | atcaataagcagatttgttaataaatcctttaaagacagtaaggaatattcatatgaatatcctccttag |
| 43816 wzi check | gccgcgagcgctttctatctt |
| 43816 wcaJ check | ctgcgacacgttcgcagctt |
| NTUH wzi Fwd | cggggctgaaaatggatcttgtacaatgataaaaattgccatatgaatatcctccttag |
| NTUH wcaJ Rvs | gagctcatagtgaaaatttgttaactagcacaacatataaattttggactataaatgcgtgtaggctggagctgcttc |
| NTUH wcaJ check | ttcggtcttttcacgggagc |
| NTUH wzi check | cattgtgagctgggtgggtg |
| pWKS130 inverse F4 | gctggcgtaatagcgaag |
| K1 wcaJ gRNA 1 F | gtggatgtggtcgaggtcctacaagttttagagctagaaatagcaagtt |
| K1 wcaJ gRNA 1 R | cttgtaggacctcgaccacatccacacattatacgagccgat |
| pCRISPR inverse R3 | tacgtagcagattgtactgagagtgc |
| pCRISPR-K1 wcaJ 5’ HA F1 | gcactctcagtacaatctgctacgtagtaggcggtaattcagtcattacg |
| K1 wcaJ 5’ HA R1 | cctcttacaaggttgccatgaagatttcc |
| K1 wcaJ 3’ HA F1 | ggaaatcttcatggcaaccttgtaagaggttggagcatttggttaga |
| pCRISPR-K1 wcaJ 3’ HA R1 | ggtttcttagacgtcaggtgctcgagtgccgccatcaatgatgatg |
| K2044 wcaI seq F1 | gctgattgtcatcttgttgttc |
| K2044 gnd seq R1 | cagaatcggagcaactaactc |
| AR0361 waaL gRNA 2 F1 | gtggatcaccagcatgatgaccacgttttagagctagaaatagcaagtt |
| AR0361 waaL gRNA 2 R1 | cgtggtcatcatgctggtgatccacacattatacgagccgat |
| pCRISPR-waaL 5’ HA F1 | ggtgcactctcagtacaatctgctacgtaccggatattttgctgttcac |
| AR0361 waaL 5’ HA R1 | catcccactagtgtctggcgaagttactcac |
| AR0361 waaL 3’ HA F1 | cttcgccagacactagtgggatgtttatcatcagcaatc |
| AR0361 waaL 3’ HA R1 | ggtttcttagacgtcaggtgctcgagcctccgaggataaagacttct |
| AR0361 waaL seq F1 | ctctataccatcgtcgacatg |
| AR0361 waaL seq R1 | gagaggtcgaactgcgtatc |

**Supplemental Table 2.** Primers used in this study.
